# Honey contamination from plant protection products approved for cocoa cultivation: a systematic review of existing research and methods

**DOI:** 10.1101/2022.12.26.521958

**Authors:** Richard G. Boakye, Dara A. Stanley, Blanaid White

## Abstract

Cocoa (*Theobroma cocoa*), which is the key ingredient of chocolate, is an important economic crop plant which supports the livelihoods of an estimated forty to fifty million people directly involved in its cultivation. Many cocoa producing countries, especially those from the developing world, rely on the income from cocoa export to support their economies. The plant is, however, prone to disease and pest attacks and therefore requires the application of large volumes of pesticides to guarantee satisfactory productions. Even though pesticides help protect the cocoa plant from disease and pest attacks, unintended effects of environmental contamination are also a possibility. Honey, a product of nectar collected by honeybees from flowers during foraging, may be a useful proxy for the extent to which landscapes are exposed to pesticides and the degree of pesticide accumulation in the environment. The overreaching question is: to what extent has the effect of pesticides imputed for cocoa production on honey received attention in research? In this present study, we conducted a systematic approach to quantify existing studies on honey contamination from plant protection products approved for cocoa cultivation. We observed that one hundred and sixty-nine different compounds, comprising some recommended and other unapproved compounds for cocoa cultivation, were detected in 81% of the reviewed 104 publications. Our results further point to the neonicotinoids as the most detected class of pesticides, with imidacloprid particularly being the single most detected compound. However, the most remarkable observation made from this study points to disproportionate studies of honey contamination from pesticides conducted in cocoa and non-cocoa producing countries with only 19% of the publications taking place in the latter. To bridge the gap, we suggest prioritising increased research in cocoa growing countries to ameliorate the significant gaps in knowledge owing to limited studies emanating from these geographic regions.

## 1.0 Introduction

Cocoa (*Theobroma cocoa*) is an essential economic crop with widespread global demand and uses owing to its rich protein, carbohydrate, fats and vitamin contents [1]. An estimated 40 to 50 million people depend on cocoa cultivation for their income [2]. Many cocoa producing countries, especially those from the developing world, heavily rely on the income generated from its export to support their economies [3, 4]. The economic value of cocoa is estimated in the region of US$11.8 billion from global annual production of 4.2 million metric tons of cocoa beans [2]. The overall cocoa confectionary market generates about US$80 billion worldwide [5] with West Africa being its main production hub. The cocoa plant is, however, vulnerable [6, 7] to attacks from the cocoa swollen shoot virus, beetles and capsids (miridae) and phytophthora pod rot (commonly called black pod) [8, 9]. Between 20% to 30% of cocoa produced around the world is lost to pest and disease attacks [10]. The black pot disease alone accounts for the loss of 700,000 metric tons on global scale [11]. An economic loss to the tune of US$3.16 billion from production losses caused by insects pests and diseases has been reported [12]. The potential annual production levels are therefore hampered by the negative effects of pest and disease.

In view of the impact of disease and pests on cocoa plants, concerns have been raised about possible threats to annual production even though efforts are being made to ensure increased outputs [13]. Pesticide application has therefore become a highly popular approach for its cultivation to prevent losses and help meet global demands [14, 15]. In Nigeria alone, an estimated 125,000 to 130,000 metric tons of pesticides are inputted for cocoa cultivation in order to safeguard production levels [16]. Even though it is acknowledged that there has been a more focused use of pesticides on cocoa farms, concerns have still been raised about environmental pollution from the use of pesticides for cocoa production [16]. Some of these concerns are related to the potential impact of pesticides imputed for cocoa cultivation on the general environment and other food substances produced nearby [17]. This is premised on the fact that a significant proportion of applied pesticides also find their way into the environment [18]. Just about 0.01% of applied pesticides are determined to reach their target but the rest filters into general ecosystem [18, 19]. The use of pesticides therefore holds inherent potential to contaminate the environment and threaten human health at some level of concentrations if they enter into the food chain [20, 21].

Honey, which is a sticky scented food, is a transformed product of nectar and other emissions from flowers and plants, produced by honeybees and stingless bees [22, 23]. It is a highly concentrated solution consisting largely of sugars enriched with protein, mineral, vitamins, organic acids, amino acids and polyphenols as the key micro-nutrients [22, 24]. Honey is recognised and marketed as a functional food owing to its intrinsic health-promoting properties from clinical confirmations [25]. As a source of sugar, honey has been used in the production of honey wines and beer and as additives in breakfast cereals and bakery goods [26]. Beyond its value as food, its application has essential medicinal values [23]. It is been found to be a rich source of antioxidants which have been identified as having potential in the treatment of different ailments including cancer, cardiovascular diseases, inflammatory disorders and upper respiratory infections [23, 27]. Saha [28] indicated that honey has been proposed for the treatment of gastrointestinal, heart-related and inflammatory conditions. The presence of phenolic acids, flavonoids, ascorbic acids, proteins, carotenoids and enzymes like glucose oxidase and catalase in honey are identified as potential sources for the proposed health properties of honey [29].

The maximum residue limit (MRL) is the highest permissible level of pesticide residue (expressed as mg/kg) recommended by the Codex Alimentarius Commission as legally accepted in food commodities and animal feeds [30]. MRLs serve as guidelines and codes of practices to safeguard the health of consumers and to guarantee fairness in food trade commodity [31, 32]. Created in 1963 by the FAO as an international body [33], the Codex Alimentarius is mandated to develop safety requirements for all food and animal feed and related texts which are presented in a uniform manner [30]. The European Union takes into consideration the MRLs specified by Codex Alimentarius as well as issues of good agricultural practices in setting MRL within its regional block [32]. On 23^rd^ February, 2005, the European Parliament and Council adopted Regulation No. 396/2005 (which amended the Council Directive 91/414/EEC) which articulated MRLs of pesticides in food and feed from plant and animal sources [34] within the European Union which is one of the world’s biggest market for agricultural products. The model for calculating the MRL was revised by EFSA to streamline its calculation and ensure conformity with the internationally agreed assessment methodology following Joint Meetings on Pesticide Residues [35]. As a natural food produced by *Appis mellifera,* honey is deemed a food substance of animal origin under Directive 2001/110/EC and therefore needs to meet specified requirements. The EU has established MRLs for pesticides in honey which range from 0.05 mg/kg to 0.2 mg/kg [36]. Where an MRL is not specified, the default limit of 0.05 mg/kg is applied in honey [37]. However, different national or regional bodies also tend to set different upper pesticide residue concentration limits but this has been cited as a source of confusion in international markets [38]. Harmonization and standardisation of MRLs is therefore a necessity.

Honey bees collect water, nectar pollen and other resources from the environment into the beehive for the production of honey [39]. The foraging behaviour of bees throughout the general environment means that bees can transport foreign bodies which may compromise colony products. Honey, as a hive product, is susceptible to potential contamination from the use of applied pesticides for crop production [40] and can be used as a proxy to evaluate the general state and health of the environment because of its ability to reveal the chemical condition of the environment [41, 42]. The levels of contaminants in honeybees, honey and pollen have been used to assess the level of heavy metals in the environment in both natural and human disturbed landscapes [43, 44]. Similarly, Perugini et al., [45] used contaminations in honey bees, honey and pollen to access the presence of polycyclic aromatic hydrocarbons in the environment. It is clear therefore that studies of pesticide contamination in bee products such as honey can be used to assess the extent of pesticide use in a particular environment. This review utilises this relationship between pesticide contamination of honey and pesticide use in the broader environment to evaluate the current state of knowledge of the extent to which pesticides used for cocoa cultivation have been detected in honeys in different geographical regions. Specifically, the study seeks to:

- Examine the period and geographical spread of existing studies of contamination of honey by pesticides approved for cocoa cultivation.
- Synthesise the different classes and types of pesticides studied during the study period in the different regions.
- Assess the concentrations of detected pesticides in relation to the maximum residue limit (MRL) set by the European Union
- Evaluate the different approaches and techniques usually applied for the extraction and analysis of pesticide residues in studies.

## 2. Materials and methods

### 2.1 Formulation of search strings

A set of set specific search strings were developed in relation to the objectives outlined for this study. Prior to developing the search strings, a preliminary study was undertaken in Google Scholar to identify key publications that report on the pesticides recommended for cocoa production to guide the construction of the search strings. Eight publications (seven peer reviews and one international report) were found that reported on approved pesticides for cocoa cultivation in key cocoa producing countries, namely Ghana, Nigeria, Cameroun and Ivory Coast, which account for 70% of the world’s cocoa production [46]. These publications facilitated the compilation of a list of the key pesticides comprising 23 insecticides, 17 fungicides and 2 herbicides approved for cocoa cultivation (S1 Table). Following the successful compilation of this list, three sets of search strings (S2 Table) were formulated for literature search. Search strings were broken into two to reduce the length of the string for each category of search engine.

### 2.2 Literature search

The search strings formulated were used to search within Web of Science Core Collection, PubMed, and Scopus to retrieve publications which satisfied the inclusion criteria set out for this study. The Web of Science Core Collection and PubMed were searched on 12^th^ October 2020 following which 891 and 469 papers were retrieved respectively which were exported to EndNote. The output of 1,360 datasets retrieved from these two search engines were strictly peer reviewed publications and therefore books or book sections, reviewed papers, theses, and grey literature were not exported. Grey literature were excluded from the study as they have been found to sometimes exhibit potential lack of research method stringency [47]. Further to this, only journal articles published in English were searched for the study due to limitations in speaking and writing in non-English foreign languages. Even though searches using multiple languages would have been more advantageous, using only English did not offer any considerable bias because study has established that 90% of natural science research is published in English [48]. A further search was conducted in Scopus on 26^th^ October 2020 using the developed search strings which resulted in 524 retrieved articles which were also exported into Endnotes bringing the total dataset of retrieved article to 1,884. The author lists and titles of exported dataset were compiled and listed in Microsoft Excel to identify and remove duplicates. Duplicates based on the combined list were then removed using conditional formatting in Excel resulting in a dataset of 1,282 articles. These were then screened based on the titles and the abstracts to assess the studies which report residues of active ingredients of pesticides in honey. However, no restriction on study design, date, or geographical zones were applied during the literature search for the relevant studies for this review. The screened data yielded a total of 91 papers. However, one article was not accessible. An email was sent to the corresponding author with a request for the paper, but no positive response was received. Subsequently this paper too was excluded. The shortlisted papers for inclusion at the beginning of the study comprised 90 articles. These were subjected to quality assessment prior to data extraction. This resulted in the further exclusion of 2 studies (see section 2.2). The flow chart in (Fig 1), based on [49] shows the procedure taken to arrive at the included publications at the start of this study. However, at the time of submitting the manuscript, twenty-four months had elapsed. Therefore, an up-to-date data search was performed to retrieve probable studies published from November 2020 to November 2022. The three databases being used for this study, namely Scopus, PubMed and Web of Science were subsequently searched on 27^th^ November 2022 using the search strings developed for this study. The search resulted in an initial list of 2,610 publications. Refining it through automation by limiting the years from November 2020 to November 2022 and further limiting it to only peer reviewed English published journals reduced the number to 261 (Web of Science=111, PubMed=142 and Scopus=16). These were exported to EndNote which was then used to remove duplicates. This was followed with a manual check for further removal of any duplicates still existing. Total duplicates removed using EndNote and a manual check was 78. The abstract and titles of the remaining 193 publications were then screened following which 170 were excluded as having not studied residues in honey. This resulted in 23 publications. However, the publication by [50] was not accessible and it was therefore excluded before text screening and quality assessment.

**Fig 1.**
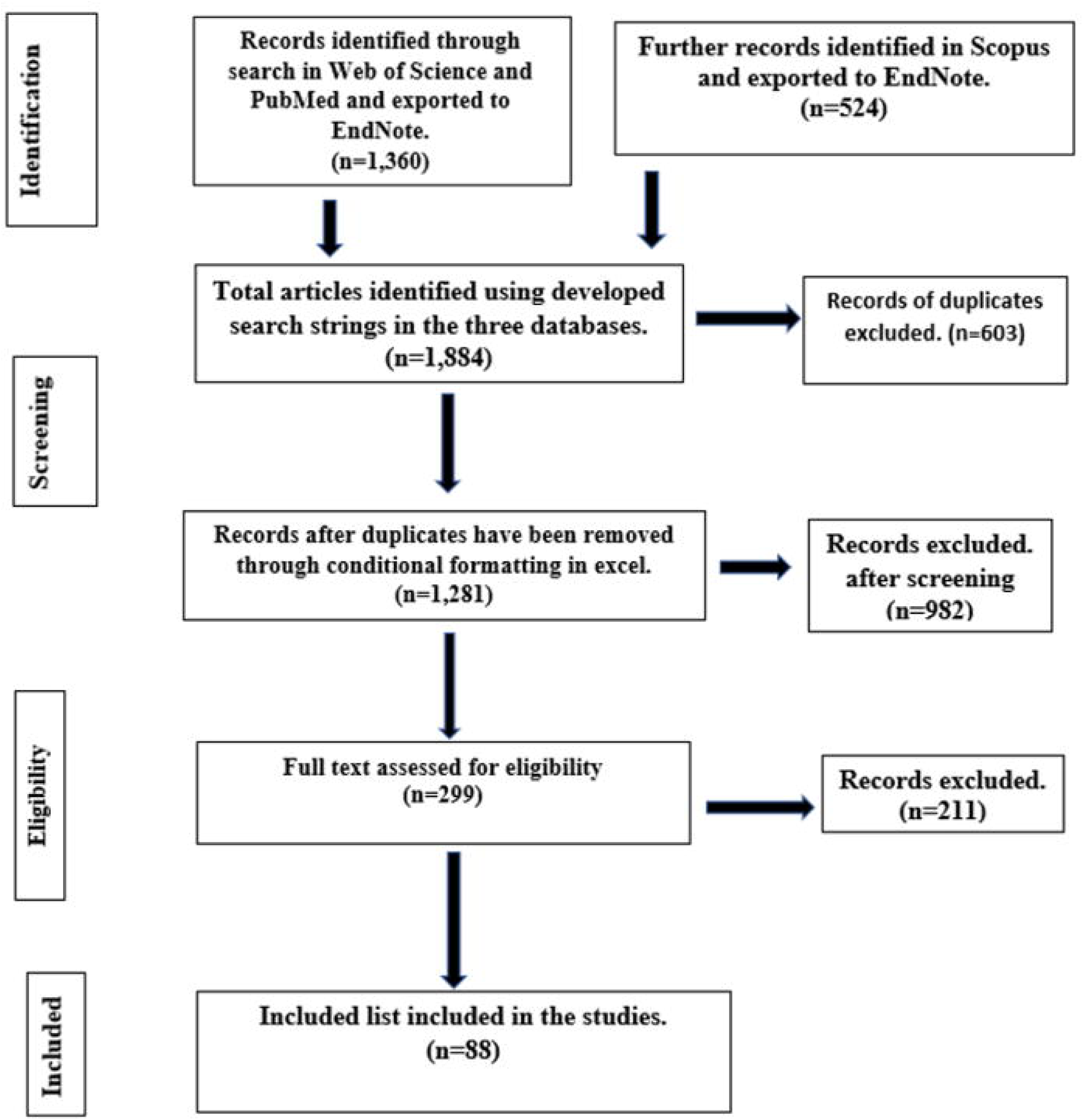
Procedure followed to select journals for inclusion in this systematic literature review. Based on PRISMA flow chart [49].

#### 2.2 Quality assessment

Before conducting data extraction, each selected study was appraised to assess it in terms of quality and suitability (S3 Table). This allowed us to evaluate the strengths and limitations of each study [51]. The appraisal process enabled us to include only studies which were of sufficient quality for this review. Our study was designed to address multiple study objectives by evaluating a broad range of multiple issues that included differences between countries of study, analytical techniques, extraction techniques, detected pesticide residues in the context of limit of detection and quantification, among others. In order to have reproducible criteria for the critical assessment of the quality of all selected studies and also taking into consideration the aims of our review, we applied a checklist of eleven customized questions (S8 Appendix) based on proposed checklists [52] for evaluating quantitative studies. The included studies were appraised by two reviewers using the scoring system previously applied by [52]. In this grading system, selected studies are scored to what extent they satisfied the criteria (“yes”=2, “partial”=1, “no”=0, NA=not applicable). The scoring systems developed by [52] had been designed using the guidelines previously developed by [53] and [54]. In our study, the first reviewer appraised each selected study which was subsequently validated by a second reviewer. The overall scores agreed by reviewers ranged from 45% to 100%. Based on the outcome of the quality assessment conducted, reviewers agreed for eight studies (two from the previous search and six from the up-to-date search) to be excluded. Of these eight studies, three analysed pesticides residues in honey samples were collected from multiple countries and therefore the study could not be assigned to one specific country for analysis. The other five studies only utilised blank honey as a sample matrix exclusively to demonstrate the robustness of the analytical method but did not quantify concentrations of pesticide residues in those samples. They were therefore deemed ineligible for further assessment. Overall, one hundred and four (104) studies were deemed to have satisfied both the inclusion criteria and quality assessment for inclusion in this review (S3 Table & S6 Fig)

### 2.3 Data extraction procedure

A comprehensive review of the full test of each included paper was conducted to identify relevant data which were subsequently extracted. The set of various data extracted from included studies, which covered information of year of publication, geographic location of study, types of pesticides studied and detected, extraction and analytical techniques, among others are captured in Table 1. Furthermore, qualitative data such as summaries of the abstracts, key findings, aims and study design of each article were also captured as part of the data extraction.

**Table 1.**
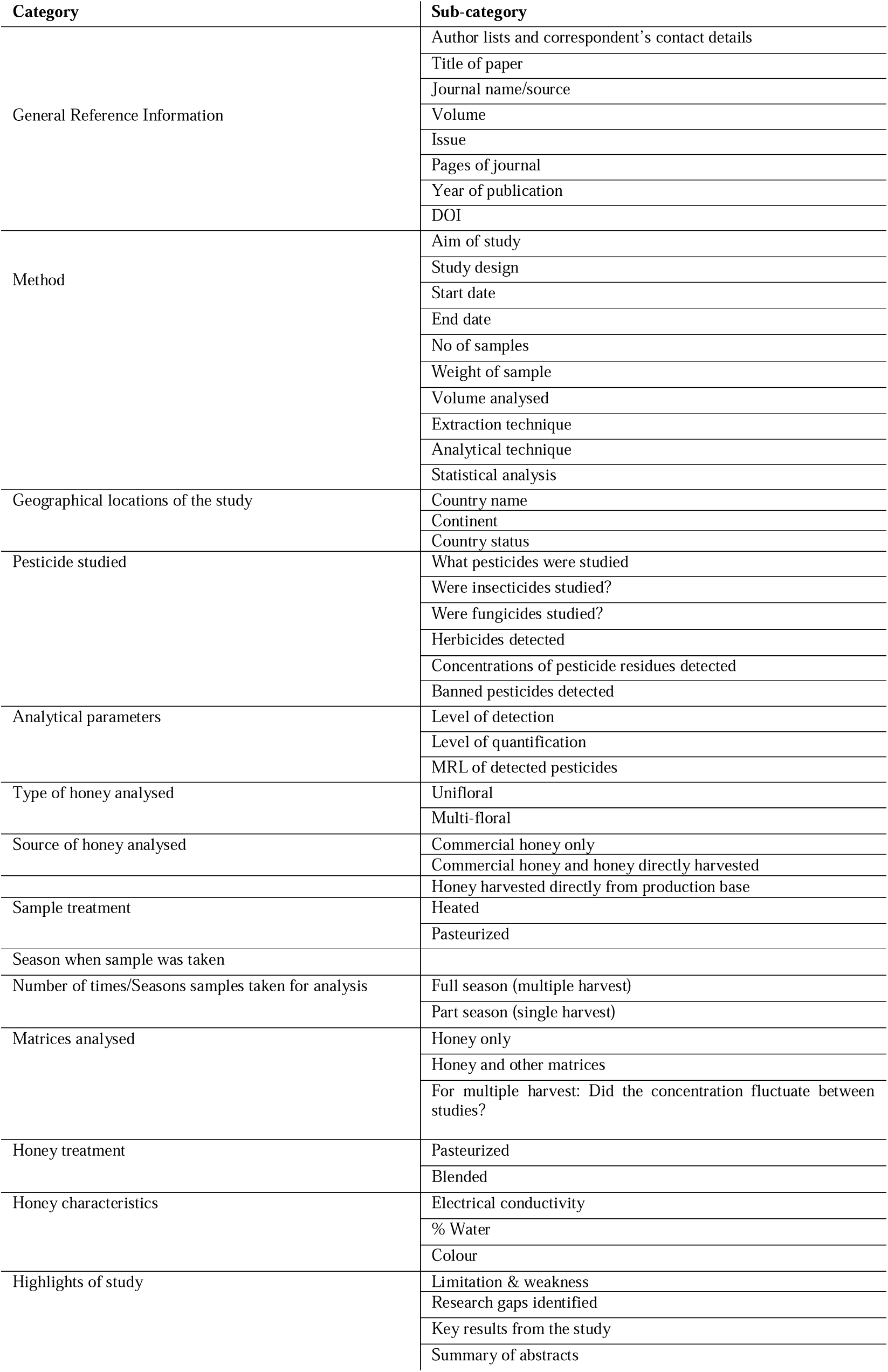
Type of data extracted from included articles. Based on this, a customised data extraction form was developed and used for data capture and processing.

Pesticides were recorded in three categories, namely insecticides, fungicides, and herbicides. Pesticides targeted in each study were recorded and those detected subsequently identified and their concentrations recorded and compared with the maximum residue limits (MRL) set by the European Union. This was carried out to determine those which exceeded the MRL which is expressed as the milligram of residue per kilogram of feed commodity (mg/kg). All units of concentrations of detected pesticides given in different units were converted to mg/kg.

## 3. Results

### 3.1 Geographical spread and period of study

The results reveal that the first of the 104 papers included in this study was published in 1997. However, publications were not sustained immediately after 1997 as the next published paper was recorded five years later (Fig 2A). Even though the trend of publication persisted thereafter with no breaks, publications started experiencing exponential growth from year 2015. Seventy-three percent of the included studies were published from 2015 to 2022 with at least 7 publications per year except the year 2021 where 4 studies were published. Overall, the year with the highest number of studies was 2018 which accounted for 15% of the included studies (Fig 2A & B). The aim and key findings from each study are summarised in the data set (S4 Table).

**Fig 2.**
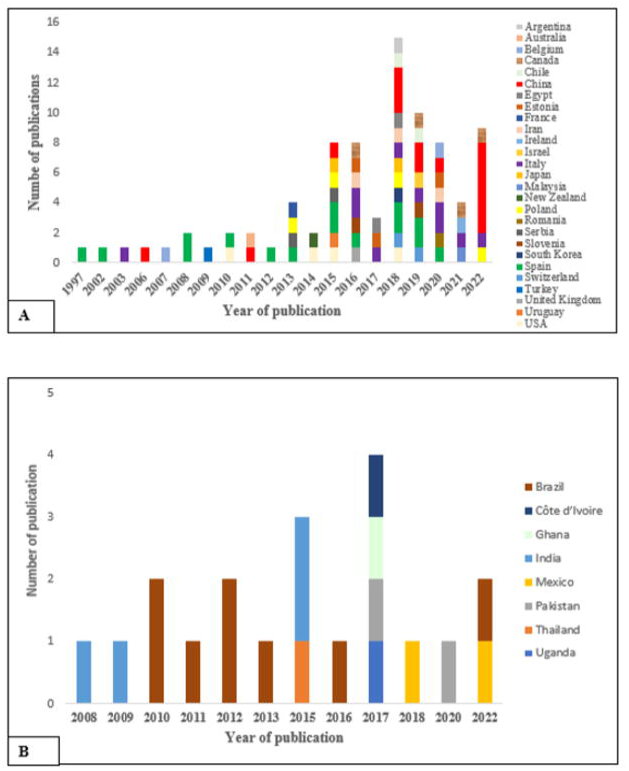
The year and country where existing studies were conducted. The studies were published over a 25-year period across 35 countries. A) Non-cocoa growing countries where studies were conducted with the first study taking place at Spain. B) Cocoa producing countries where studies took place. Overall, 20 studies were conducted in eight cocoa growing countries with the first study taking in India eleven years after the first study was conducted in Spain.

Our findings show that the spread of the studies was skewed towards Europe (43%) and Asia (30%). But even though the highest number of the studies took place in Europe, they were concentrated in Spain where 33% of the 47 studies conducted in Europe took place. Similarly, 48% of the 31 studies conducted in Asia took place in China. Overall, the highest proportion (81%) of the included studies took place in twenty-seven countries where cocoa is not cultivated. Further observation made revealed that at least one study was conducted on each continent except Antarctica where no studies took place (Fig 3).

**Fig 3.**
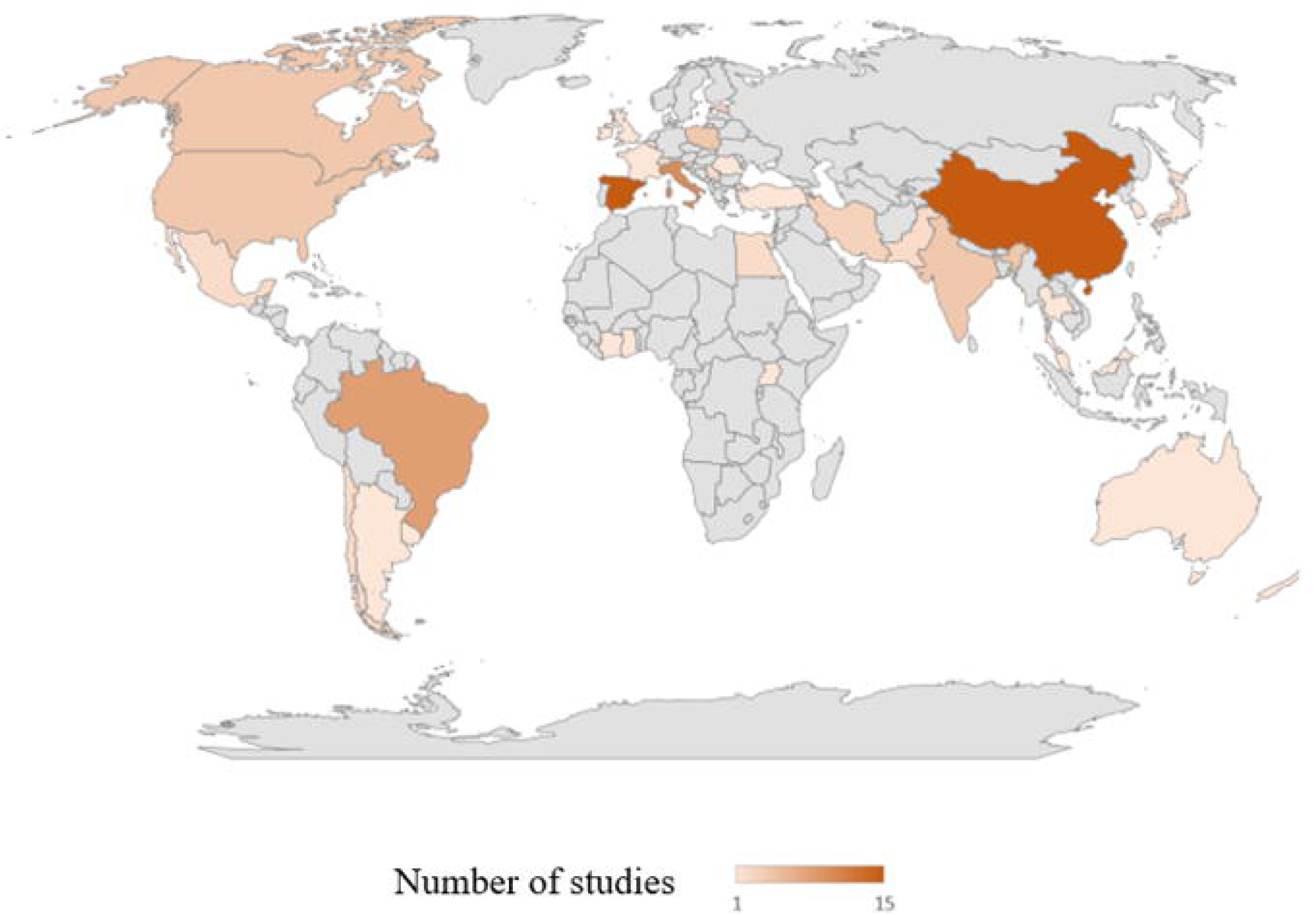
Geographical spread of the studies that were undertaken (grey areas=no studies). Spain (15 studies), China (15 studies) and Italy (10 studies) were the top three countries with most studies. One study each was conducted in Ivory Coast and Ghana which are the first and second ranked cocoa producing counties in the World respectfully.

Out of the 20 studies which took place in cocoa producing countries results show that 8 studies were conducted in Brazil which is ranked sixth highest cocoa producing country in the world and accounts for 5% of global production of cocoa beans (S5 Table). A further 4 studies were carried out in India ranked as the sixteenth cocoa producing country with production levels being less than 1% of global production. Only one study was conducted in each of Ivory Coast and Ghana which are ranked first with 39.1% production and second with 17.0 % production respectfully in global cocoa productions. Mexico (2 studies), Pakistan (2 studies), Thailand (1 study), and Uganda (1 study) make up the rest of cocoa producing countries where studies took place. The eight cocoa producing countries where studies took place represent 14% of the 57 cocoa producing countries worldwide (Fig 4).

**Fig 4.**
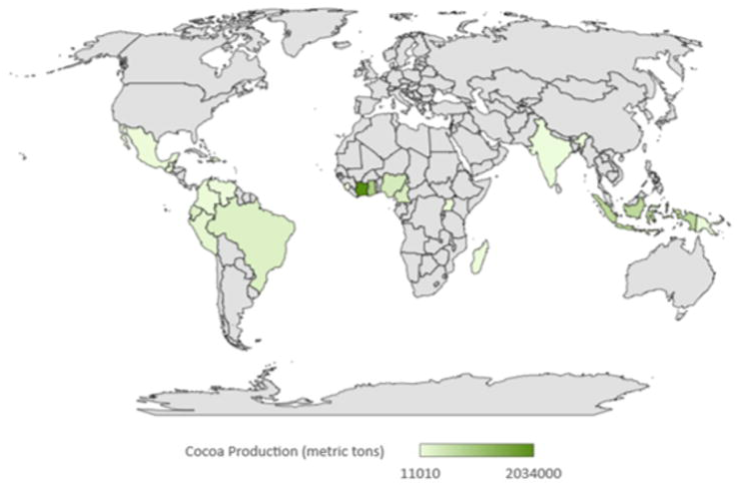
The 57 cocoa producing countries in the world based on the metric tons of cocoa produced annually. Ivory Coast is ranked first with 2,034,000 metric tons of annual production. 19% of the included studies took place in eight cocoa producing countries. (Grey areas = non-cocoa producing areas).

### 3.2 Limit of detection and limit of quantification applied in studies

The Limit of detection (LOD) is the validated lowest concentration of a trace substance detected using regulated or standard procedures [55, 56] and the limit of quantification (LOQ) is the smallest concentration of a substance which can successfully be quantified [57, 58]. Both LOD and LOQ were evaluated for all pesticide residues that were detected in the included studies (S3 Table & S4 Table). Our result showed that 80% of included studies applied LODs which were below the specified EU MRLs for the studied compounds. However little information was provided in 18 studies to establish the applied LODs. From the studies with clearly stated LODS, it was only in three studies where the reported LOD exceeded the MRLs. Similarly, most studies (77%) clearly indicated the applied LOQs which were found to be lower than the MRL. In 17 others, limited information was provided on applied LOQ. From the clearly stated LOQs, three were found to exceed the MRL. Furthermore, in two other studies, the LOQ did not exceed the MRL, but they were not sufficiently low enough for the quantification of trace elements.

### 3.3 The classes and types of pesticides evaluated

A broad spectrum of pesticides, including different types of insecticides, fungicides, herbicides other types of pesticides were analysed in honey over the years. However, our results show that insecticides received the highest attention having been studied in 91% of the eligible papers. Overall, only 4 publications studied insecticides, fungicides, and herbicides together in the same study. It was observed that pesticide residues were detected in 80% of the 104 studies reviewed. Our study found that a total of 169 different compounds, comprising of some of those recommended as well those not approved for cocoa cultivation, were detected in 86 studies which took place in 30 out of the 35 countries where studies were conducted (Fig 5).

**Fig 5.**
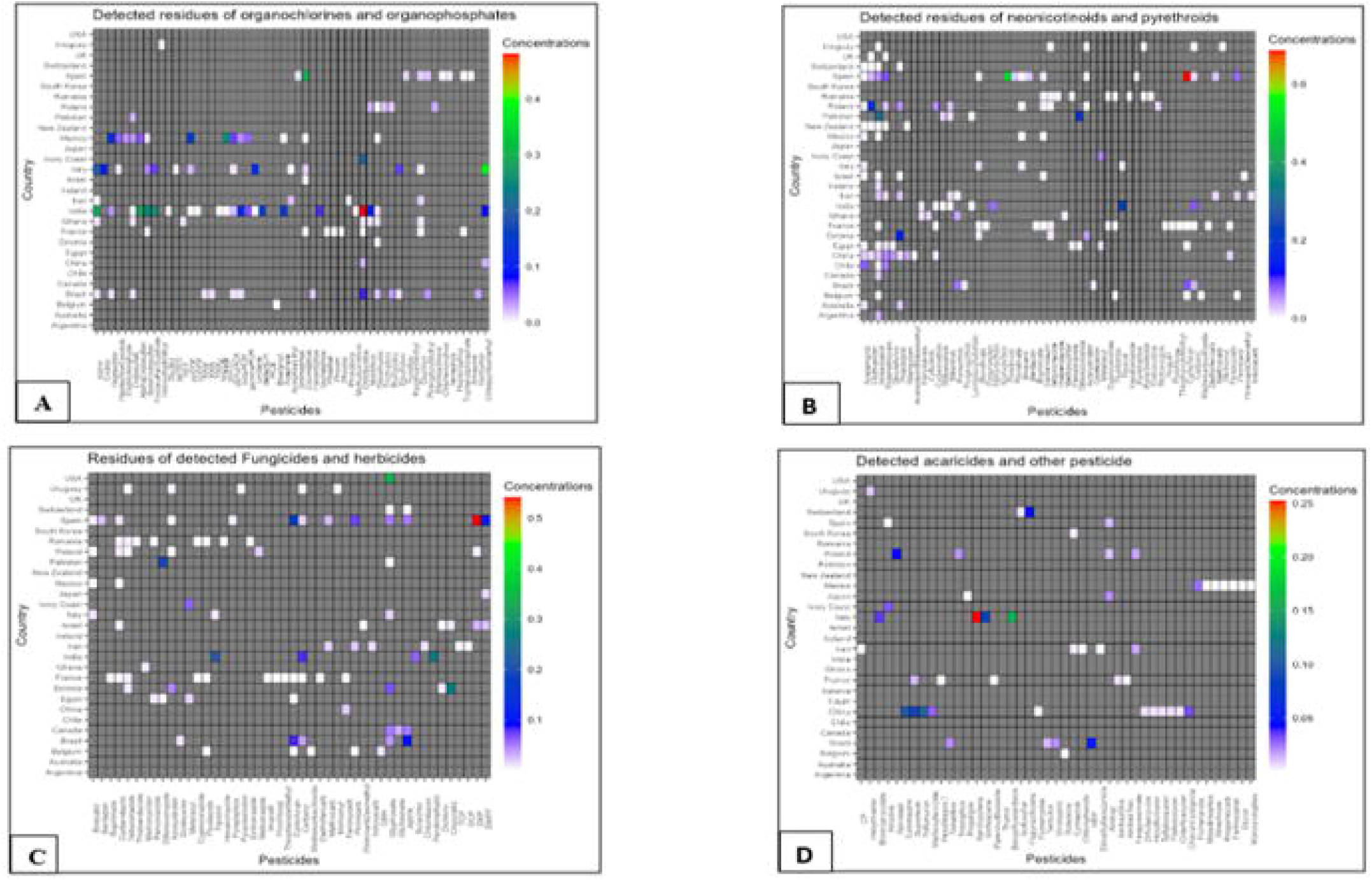
Heatmaps with colour scale on the right of each graph showing the detected: A) organochlorines and organophosphates B) neonicotinoids and pyrethroids; C) fungicides and herbicides and D) acaricides other pesticides which were studied (x-axis) over the period and the respective countries where detections took place (y-axis). To this figure, concentrations of each detected pesticide were averaged per the number of detections per country to get one value for each pesticide detected. Units for all detected pesticides were standardised by converting to mg/kg which is the unit used by the European Union which was adopted for this study. The individual graphs can be referred to at S1 Fig to S4 Fig.

The most detected classes of pesticide were the neonicotinoids with imidacloprid (detected in 20 studies), thiamethoxam (in 14 studies), acetamiprid (in 13 studies) and clothianidin (detected in 9 studies) being the most detected. Interestingly however, we observed that the first top three most detected classes of pesticides in the six cocoa growing countries, were the organophosphates, organophosphates and pyrethroids in that order (Fig 6 & S6 Table). Eleven approved insecticides for cocoa cultivation namely capsaicin, chlorantraniliprole, thiamethoxam, acetaprimid, etofenprox, indoxacarb, pirimiphosmethyl, promecarb, pyrethrum, sulfoxaflor, and teflubenzuron and one herbicide (i.e., paraquat) were not detected in any of the studies conducted in the cocoa growing countries. Additionally, our findings showed that only 2 of the 18 recommended fungicides for cocoa production (S1 Table), namely Metalaxyl-M and its isomer Metalaxyl, were detected in studies conducted in cocoa growing countries. One outcome emanating from our study shows the detection of 49 pesticides largely detected in India, Mexico and Brazil, which are not recommended for cocoa production [59].

**Fig 6.**
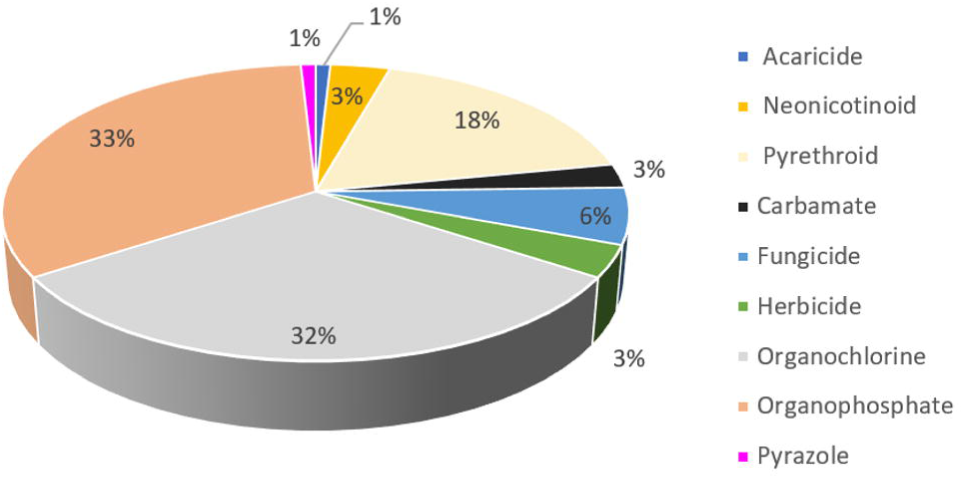
The classes of pesticide residues detected in studies which were conducted in six cocoa growing countries namely Ghana (1 study), Ivory Coast (1 study), Brazil (4 studies); Mexico (1 study); India (4 studies) and Pakistan (2 studies) where pesticide residues were detected. The highest number of pesticides residues (40 detections) were recorded in India. No pesticide residues were detected in studies conducted at Thailand and Uganda the other lists which make up for the cocoa growing countries.

The outcome from this study shows that multiple studies took place in some countries (i.e., 54% of included 35 countries) with one study taking place in the rest. Spain and China, accounted for the highest number with fifteen studies each. Among the cocoa producing countries, our results show that Brazil, India, and Mexico were the only the countries where more than one study was conducted. Our results show that some pesticides were separately detected in different studies within in the same country (seven countries in all). An assessment of the concentrations of these pesticides, where these pesticides were detected in the same jurisdiction, sowed that they were usually found to be at varying concentrations (Table 2).

**Table 2.**
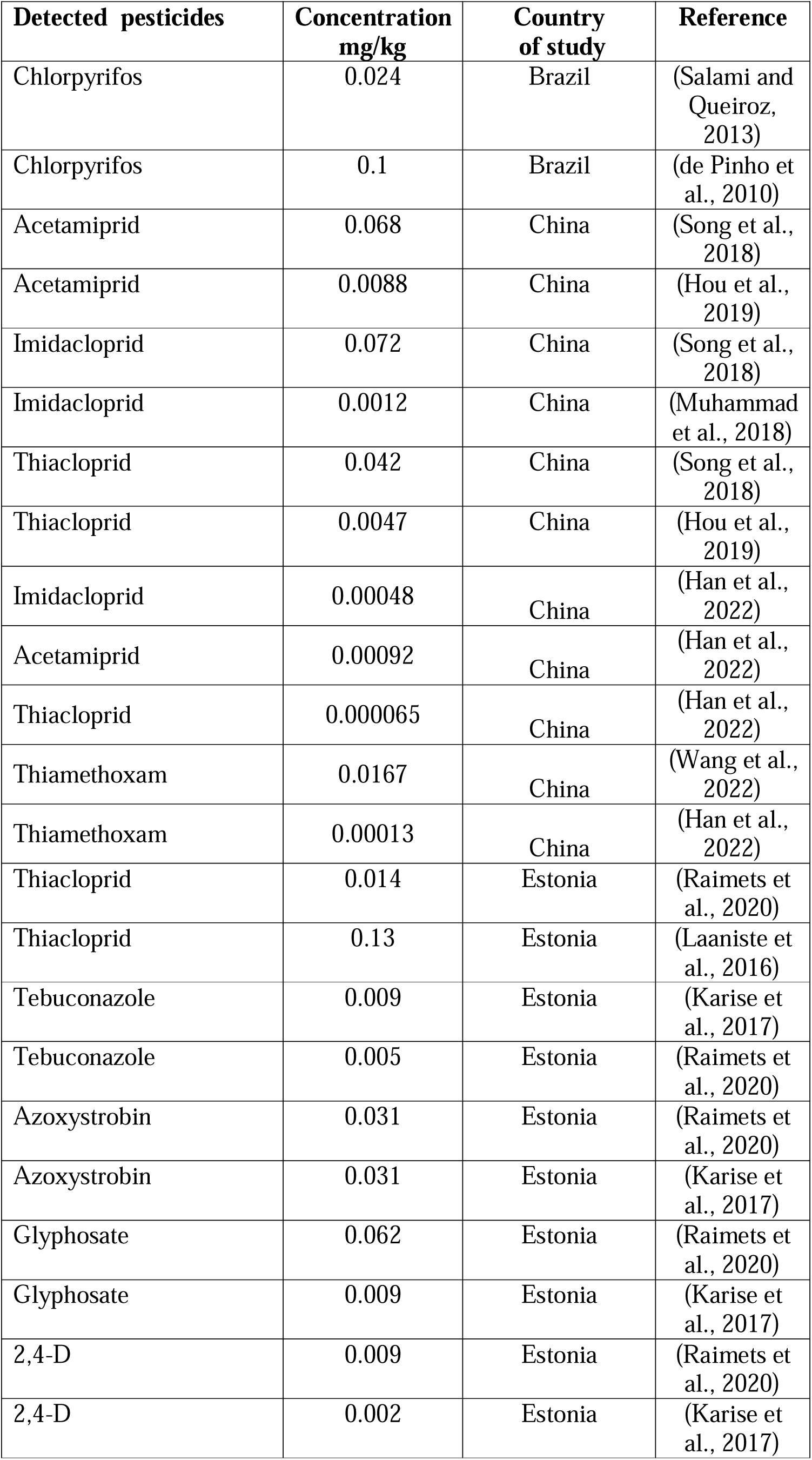
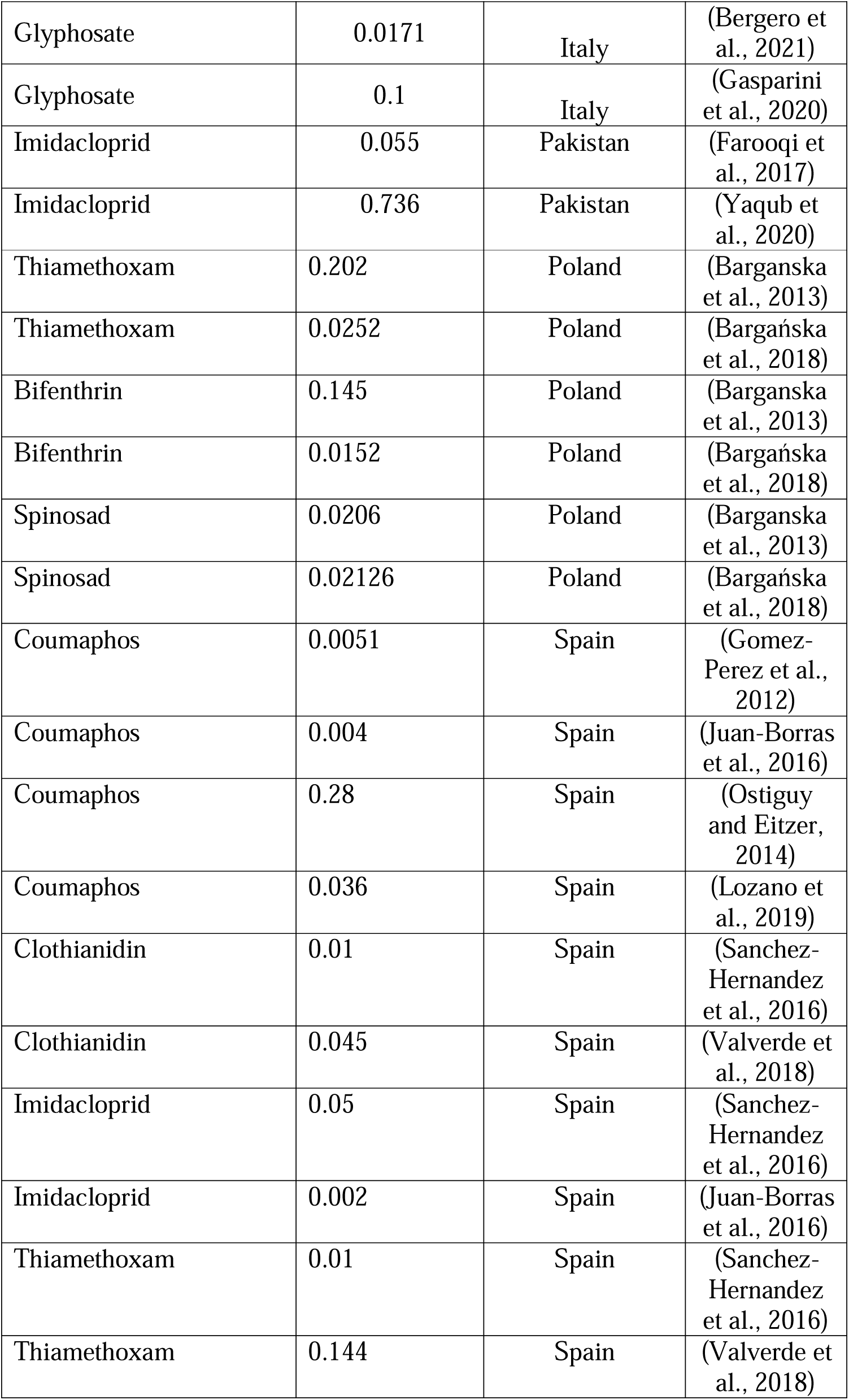
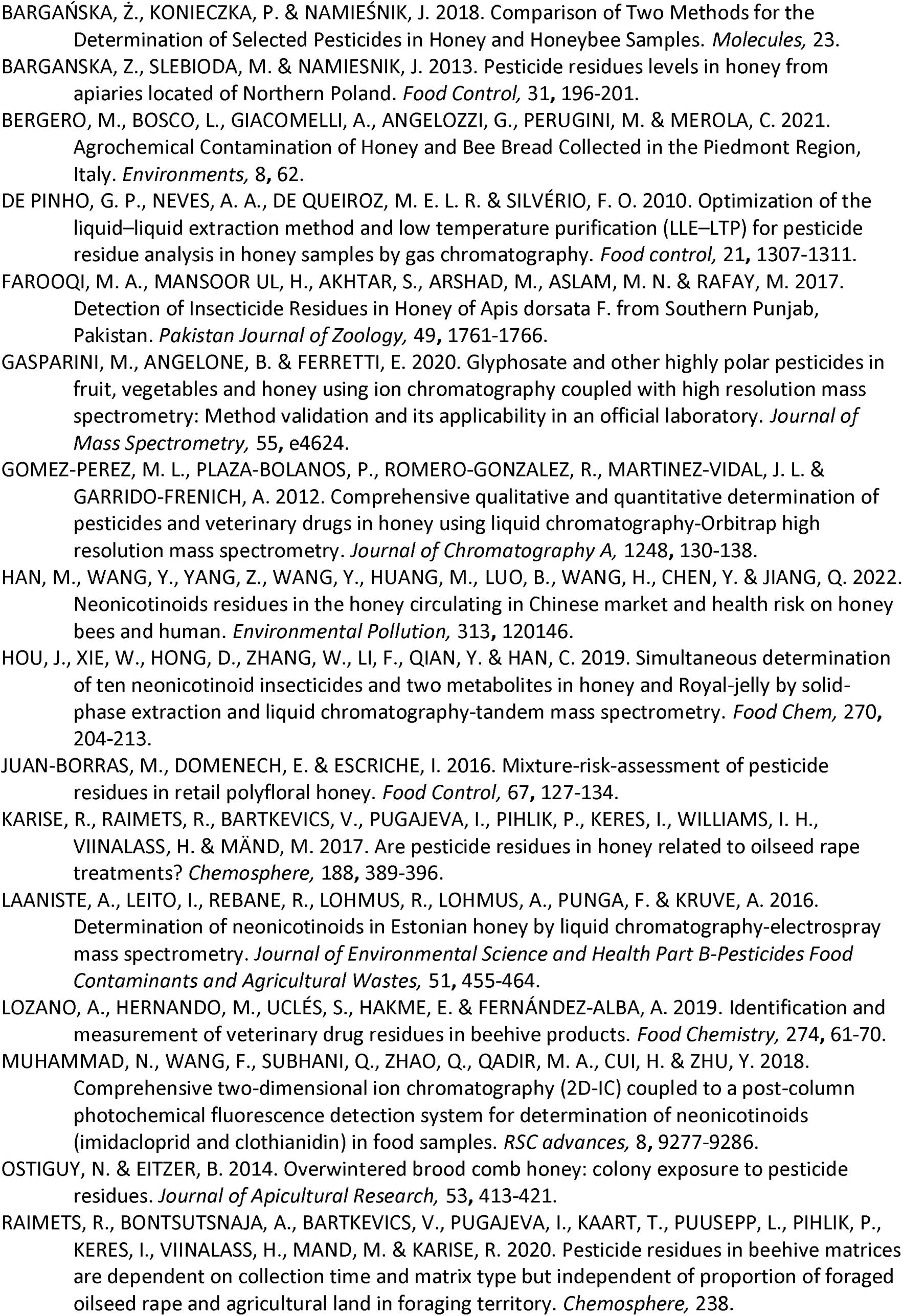
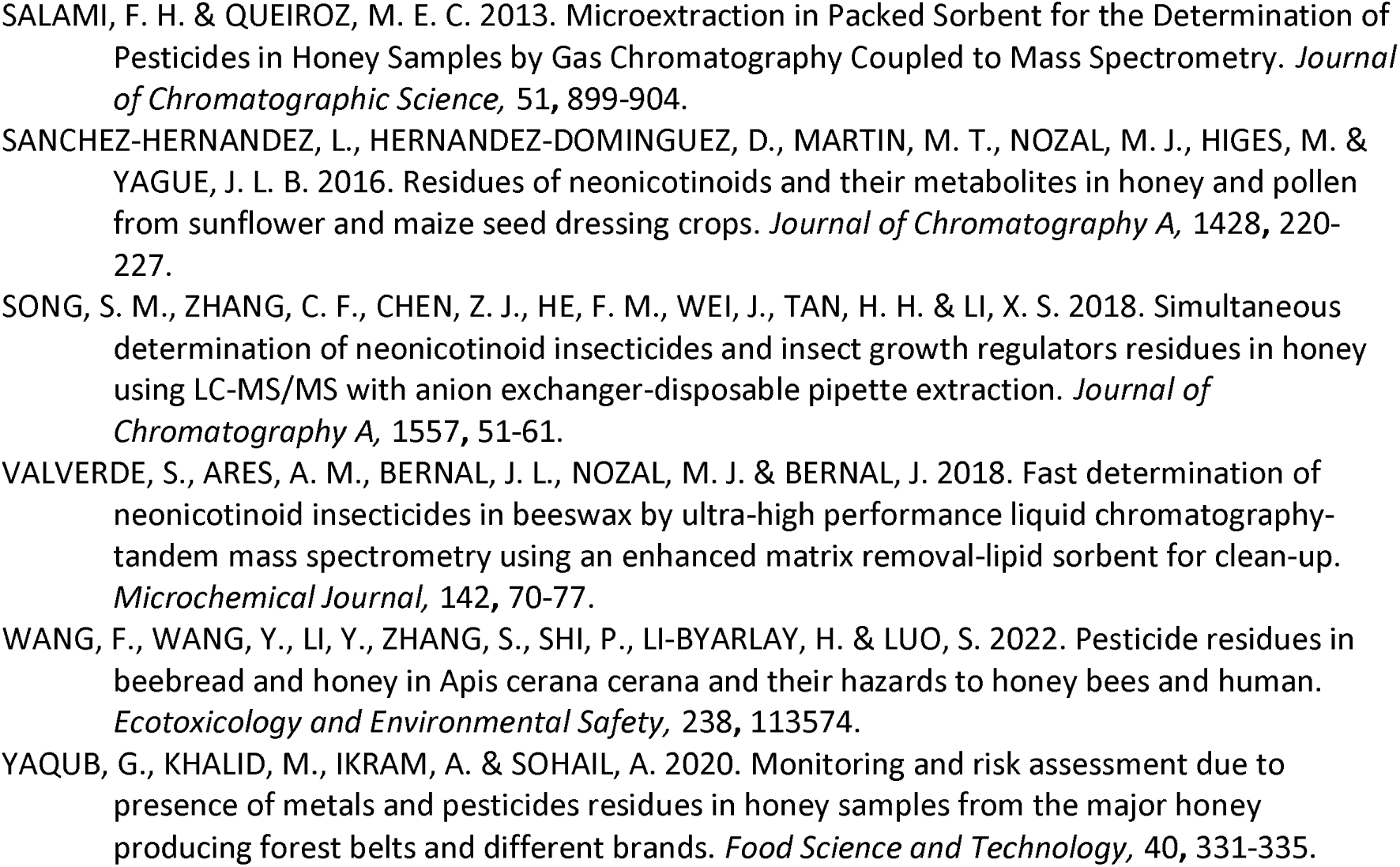
Pesticides detected in different studies within country where more than one study was conducted. Concentrations generally fluctuated except for azoxystrobin which was detected at the same concentration in two separate studies conducted in Estonia.

#### 3.3.1 Detected banned pesticides

Some pesticides are banned for use in various jurisdictions based on different legal instruments. For this study, “banned” pesticides evaluated in this study are based on the Stockholm Convention. Under the Stockholm Convention on Persistent Organic Pollutants (POs), which is signed unto by 184 states and the European Union, the production and agricultural uses of many organochlorines have been phased out or restricted because of their persistence in the environment and the danger they pose to human health [60]. All the countries where the included studies took place are parties to the convention, even though the USA and Italy are yet to ratify it. No banned pesticides were detected in 96 % of the included studies reviewed in this study. The banned pesticides were organochlorines which confirmed in three cocoa producing countries namely Ghana, India, and Mexico. In the study conducted in Ghana, Dichlorodiphenyltrichloroethane (DDT), an organochlorine insecticide which is the first of the modern synthetic insecticide manufactured primarily to fight malaria, typhus and for agricultural uses [61], was confirmed in 0.01 mg/kg concentrations. We observed that in a study conducted at Mexico, Ruiz-Toledo, Vandame [62] confirmed the presence of 10 organochlorines including heptachlor (0.13173 mg/kg); hexachlorocyclohexane (HCF, 0.654 mg/kg), endrin aldehyde (0.03564 mg/kg), and dichlorodiphenyldichloroethylene (DDE, 0.154358 mg/kg) were detected in honey from the Chiapas vicinity where official approval for their usage was withdrawn in the 2000’s. Hexachlorocyclohexane (HCH) which is used as an insecticide on fruit, vegetables and forest crops, and its isomers, endosulfan and aldrin, were detected in the studies which were conducted in India at concentrations of 0.0028 mg/kg, 0.00253 mg/kg and 0.00201 mg/kg respectively. It must however be pointed out that, unlike the other three counties where DDT is completely banned from use, DDT is still permitted for fumigation against mosquitoes as a malaria controlled measure in India [63]. In Spain, DDE (0.09-0.6598 mg/kg) a metabolite of DDT, was detected in one out of the sixteen studies which were conducted.

### 3.4 Exceedance of EU MRLs

Overall, 13% of the included studies in ten countries recorded concentrations of pesticide residues which exceeded the maximum residue limit (MRL) set by the European Union in honey (Table 3). Of note, EU MRLs are occasionally revised in light of additional scientific data becoming available to the European Food Safety Authority, and during the time period of this study, these revisions resulted in an increase in MRLs for certain pesticides, It was additionally observed that there was one study where the malathion concentration was found to have exceeded the MRL set in India, but the upper limit was found to be below that of the EU MRL at the time of publication. Among the cocoa producing counties, MRLs exceedances took place in Ivory Coast, India, and Brazil. At the continental levels, our results did not show exceedances of MRLs at North America (4 studies) and Australia (1 study).

**Table 3.**
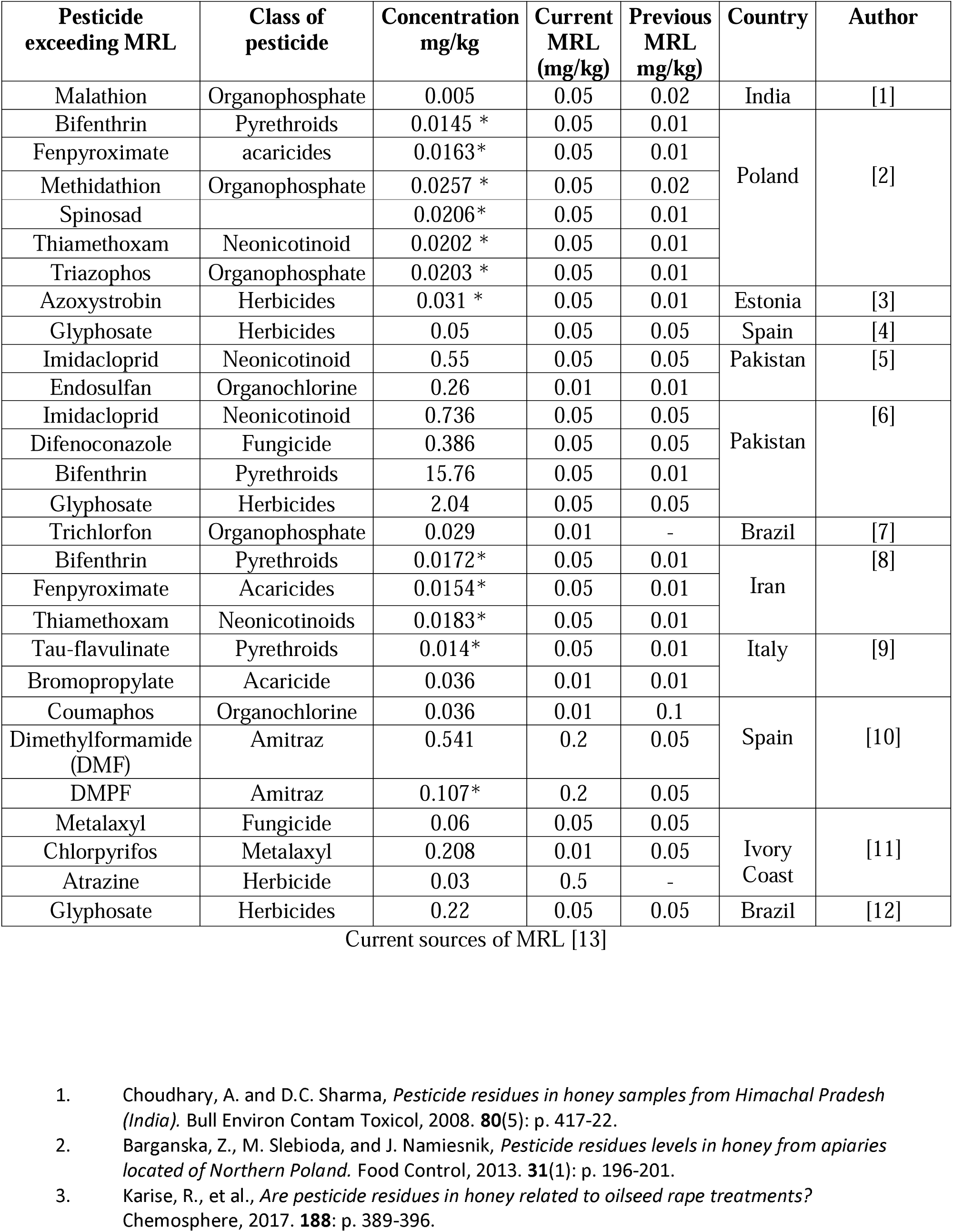
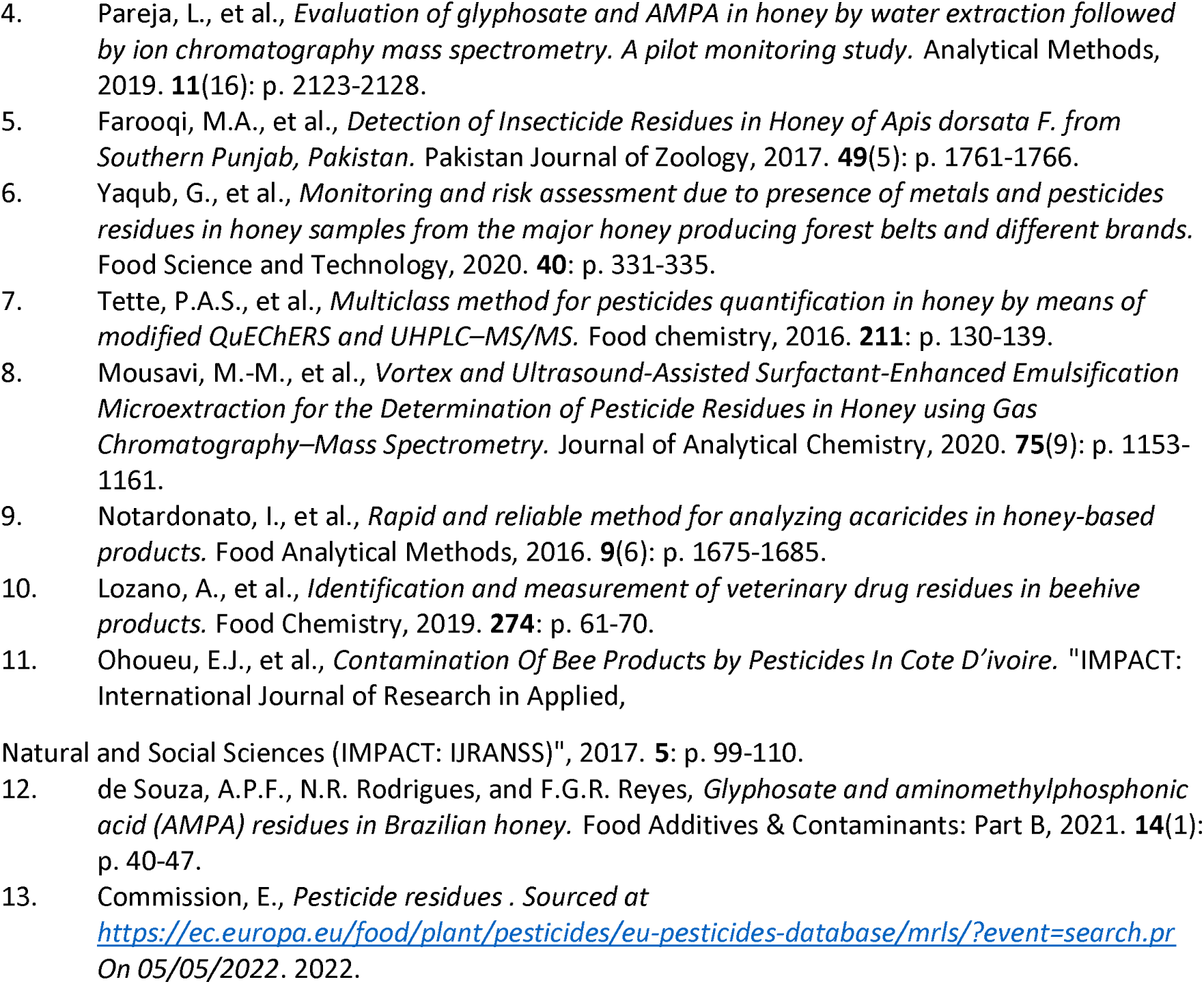
Concentrations of pesticide residues which exceeded the maximum residue limits as specified by the European Union. Concentrations marked in asterisks which were deemed to have exceeded the previous MRL set by the EU in 2005. However, these concentrations are below the revised MRL set by the EU in 2022. Malathion exceeded MRL set by India; however, the concentration recorded is below both the previous and current MRL set by the EU.

### 3.5 The types of honey investigated in existing studies

Raw honeys are unprocessed honeys which are taken directly from the beehive or from location of production [64]. However, commercial honeys differ from raw honeys in terms of processing [65] because they are processed at high temperature sometimes being heated to 70 degrees Celsius to reduce viscosity and then given rapid cooling as a way of easing handling and packaging processes [66] and to inhibit microorganism growth and reduce moisture content [67]. It was found out that both commercial honeys (35 studies) and raw honeys (61 studies) were analysed in the included studies except in one study conducted in Spain where the source of honey analysed was not specified. Both commercial and raw honeys were analysed at the same time in 7 % of the studies. Even though commercial honeys were studied over the period, limited information on honey as either being heated or pasteurised was not frequently recorded for detailed analysis. Although it has been established that heating tends to decrease honey quality with the potential to degrade pesticide residues [68], it was not possible to evaluate how these impacted the pesticide levels because information on honey either being heated or pasteurised before analysis was not frequently recorded for detailed analysis. Just 4 papers indicated that honeys were heated before analysis. None of the included studies indicated an analysis of blended honey in their studies. It was therefore not possible to evaluate parameters such as whether commercial honeys were blended. Most of the studies (61%) analysed residues in honey as the only matrice in a study. However, for the rest, residues were analysed in honey alongside other matrices which included pollen, beeswax, vegetables, honeybees, fish, beebread with the analytical technique applied also proving successful.

#### 3.5.1 Honey sampling per study

Most studies performed analysis of pesticides residues in honey samples collected on just one occasion but in 11 studies, honey samples were taken for analysis on multiple times. All the 11 studies which analysed honey samples multiple times used raw honeys. Eight of these studies collected and analysed honey samples either over a two-year period or in multiple months within the same year. Three studies analysed honey samples continuously for three years. One study uniquely analysed honey sample continuously for nine years [69]. Only one of the eleven studies, where samples were collected and assessed in multiple times across different seasons or year, took place in a cocoa growing country (Uganda).

Outcomes from such multiple sample collection recorded varied outcomes. No pesticide residues were detected in the study conducted in Uganda. It was observed that in two independent studies conducted in Chile, no pesticide residues were detected in one study, but in the other study acetamiprid, thiamethoxam, thiacloprid and imidacloprid were confirmed in three honey samples but the detections did not overly fluctuate. The other study where no trace of pesticide residues was detected in honey collected on multiple times was conducted in Spain by [70] during their two-sampling campaign in 2008. In a study conducted in France, contamination was found to be higher in samples in early spring in a study where samples were collected from apiaries in the spring, autumn, and both the early and late summer. In one study conducted in Egypt, acetamiprid and imidacloprid were detected in samples assessed both in the spring (clover season) and summer (cotton season) at different concentrations across the two seasons. One study confirmed the concentration of clopyralid and glyphosate to be higher than the MRL in Estonia a study where samples were collected and analysed in both 2013 and 2014. However, in another study in Estonia herbicides were detected in samples in collected 2013, but no sample was found to contain glyphosate in 2014. In one study that lasted nine years starting in 2004 in Estonia, an increasing trend of pesticide residues was detected throughout the years, though thiacloprid was not found to be equally distributed between 2005 and 2013 with no neonicotinoids being detected at all from 2005 to 2007. In another study, glyphosate was analysed in honey samples in 2015 and 2016 in the USA at two different sites and at both sites its concentration increased over time.

### 3.6 Extraction and analytical techniques applied

The findings from our study showed that a vast array of both traditional and other novel extraction techniques was applied for trace analysis. However, the “Quick, Easy, Cheap, Effective, Rugged and Safe” method popularly known as QuEChERS, which was developed by Anastassiades, Lehotay [71], was found to be the most prevalent technique having been applied in 41 % of in the 104 studies (S5 Fig). It was further combined with other techniques such as accelerated solvent extraction, solid phase extraction and dispersive liquid-liquid microextraction in three studies in a comparative study to assess extraction efficiencies. Our results further showed that other commonly used techniques included solid phase extraction and liquid-liquid extraction techniques which were applied for extraction in 26 % and 14 % of the studies respectively. It is also interesting to note that novel extraction techniques were utilised in 8% of studies. These included a quick polar pesticide method (QuPPe), in-coupled syringe assisted octanol-water partition microextraction (ICSAO-WPM, MEPS), the molecularly imprinted polymer (MIP)-based microneedle (MD) sensor, the dual-template molecularly imprinted polymer nanoparticles, vortex and ultrasound assisted surfactant-enhanced emulsification micro-extraction (V–UASEEME) and magnetic solid-phase extraction (MSPE). The scope of this study was limited to identifying the vast array of existing extraction techniques and their frequency of use and stretched for a comprehensive evaluation of the performances of the different extraction techniques. However, our brief assessment of the novel techniques results shows that some noteworthy successes. The QuPPe which is recommended by EU Reference Laboratories for Residues of pesticides and applied in a study in Italy achieved satisfactory validation parameters with the LOQ ranging from 0.00430 to 0.00926 mg/kg giving an indication of high method sensitivity. The ICSAO-WPM extraction technique was combined with high performance liquid chromatography and a recovery of 96.96-107.7% was achieved even enough it had low limit of detection range of 0.00025-0.0005 mg/kg. This technique involves rapid shooting of syringe which creates rapid and mass processes between phases and which in turn impacts extraction efficiency. V-UASEEME is characterised by shorter extraction time, low consumption of extraction solvents and low LOD and LOQ in the ranges 0.00003 to 0.0009 mg/kg and from 0.0001 to 0.003 mg/kg respectively. Another novel technique, the effervescence-assisted dispersion and magnetic recovery of attapulgite/polypyrrole sorbents extraction technique was developed and applied to analyse five pyrethroids in honey and it recorded limits of detection that ranged from 0.00021 and 0.00034 mg/kg with batch-to-batch repeatability of 5.05-15.01%. The technique allowed for four extraction cyclic use of the sorbent without a significant loss in the extraction recovery. A comprehensive assessment of the performances of some these different extractions have already received sufficient attention [67, 72, 73]. In summary however 48 % of the 83 studies where pesticide residues were detected applied QuEChERS for extraction. Furthermore, 46 % of the 13 studies which detected residues which exceeded the MRL applied QuEChERS for extraction. The use of graphene and carbon nanotubes, in the molecularly imprinted polymer (MIP)-based microneedle (MD) sensor, served as efficient adsorbents for dSPE clean-up which successfully removed coextractives with graphene and was found to be superior to carbon nanotubes and successfully detected chlorpyrifos in 100% of the honey samples studied. Similarly, a novel extraction technique, the Rut-MOP based magnetic solid-phase extraction, showed high extraction potential for neonicotinoids and when combined with high performance liquid chromatography was also found to be very sensitive for the detection of neonicotinoids in lemon and honey.

Just like extraction techniques, different analytical techniques were also applied for the quantification of detected pesticide residues. In all, a total of 35 analytical techniques, involving either a technique being applied independently or in combination with others, were applied for analysis. However, we observed that the most predominantly applied technique was Liquid Chromatography-tandem mass spectrometry (LC-MS/MS), which was applied in 40 % of the total studies. It was further applied in conjunction with 13 other techniques for analysis. Like the extraction, the performances of the different analytical techniques applied in the various studies were not a subject matter of this study. However, the higher frequency of use of the LC-MS/MS ultimately resulted in the LC-MS/MS detecting the highest number of pesticide residues (41% of 83 studies) and the highest number of detected residues which exceeded the MRL (4 out of 12 studies). Gas chromatography-mass spectrometry (GC-MS) was the second most frequently applied analytical technique with the technique also detecting pesticide residues which exceeded the MRL in 3 studies.

## Discussion

Honey is particularly useful to mankind both for its nutritive values and as a medium for monitoring environmental quality through the assessment of its contents for environmental contaminants. Despite being a source of both macro and micronutrients, the presence of contaminants in honey may reduce its quality thereby making it less beneficial for human consumption. Unfortunately, the probability of honey being compromised by pesticide application is a possibility because as a bee product, honey is predominantly produced in agricultural landscapes. In this present study, we undertook a systematic literature review to evaluate honey contamination from plant protection products recommended for the cultivation of cocoa (*Theobroma cacao* L.), a crop that is highly dependent on pesticides for cultivation because of its vulnerability to insect and disease attacks.

Our results show gradual but generally consistent analysis of honey contamination from pesticides residues since 1997, with peaks reporting periods taking place from year 2015. A similar observation was made by [74] who reviewed plant protection product residues in plant pollen and nectar which serves as raw material for honey. In their study they found that majority of their studies were published in the years 2012 and 2015, about the same period where most of the residues of honey started peaking as observed in our study. The increased growth after 2014 coincides with the period when the EU placed a moratorium on the use of some neonicotinoids, namely clothianidin, imidacloprid and thiamethoxam in 2013 in Europe [75, 76]. Incidentally, our results also showed that majority of the studies conducted in cocoa growing countries took place during this period. It is possible that the sharp growth in studies could be in response to the reported bee deaths due to the pervasive use of pesticides [77] and the reported worldwide decline of pollinators [78]. It is worthy of note that in 2018, a ban was placed on the outdoor use of three neonicotinoids namely clothianidin, thiamethoxam and imidacloprid [79, 80]. The possibility of the findings from the studies conducted during this period contributing to the eventual ban are likely and very much expected as the European Food Safety Authority (EFSA), which provides scientific advice for the promulgation of laws and regulations to protect consumers [81], consider published scientific research as part of their decision-making process. The year 2022 witnessed a surge in studies with most of the studies being conducted in China (Fig 2). This could be informed by the reported widespread contamination of arable lands in China to the tune of 150 million miles [82] and the reported evidence of negative effects of pesticide on public health through drinking water exposure [83].

The most remarkable observation made from this study points to a disproportionate number of studies of honey contamination from pesticides taking place in non-cocoa producing countries (81%) relative to cocoa producing countries majority of which are developing countries (Fig 3). This finding reflects the observation made by [84] and [74]. In a study which assessed the impacts on herbicides and fungicides on bees, [84] established that majority of the studies were conducted in North America, Europe and Russia. A similar trend was observed by [74] from a study which evaluated plant protection products in pollen and nectar. Cocoa thrives in hot and humid climatic conditions and tend to flourish in areas around the West Africa, East Asia and the South America [85]. Most cocoa producing countries are located outside North America and Europe. Our finding highlights a dearth of knowledge of the impact of pesticides imputed for cocoa cultivation in the environment. Considering honey as a proxy for such assessments, the paucity of knowledge may restrict a better or more detailed assessment of impacts on the general environment. Presently, the production levels of cocoa do not meet demand in several parts of the world such as China and India [86] and there is currently an increased 2.5% yearly demand of cocoa beans around the world [87]. This is likely to translate into increased production of cocoa with possible corresponding increased use of pesticides for the control of disease and insect pests. Prioritising evaluation or studies of honey contamination from pesticides application in cocoa growing areas can reveal the extent to which honey is impacted by pesticides applied for cocoa cultivation, and by extension, the extent to which these compounds are detectable in these regions.

Pesticides can be classed according to their uses and in this regard can be categorised as fungicides, weedicide/herbicides, nematicides, rodenticides and insecticides [88]. Moreover, on the basis of chemical structure, further grouping of major pesticides into organochlorines, organophosphates, carbamates, pyrethroids, triazines, and neonicotinoids also exists [42, 88]. The findings from this study provide evidence of these different classes of pesticides receiving attention in the included studies even though the compounds or classes of pesticides studied were not given equal attention across different geographic locations (Fig 5). We observed that the neonicotinoids were the most detected of all compounds (Fig S6). These findings agree with assertion made by [89], [90] and [91] that the most applied classes of pesticides around the world are the neonicotinoids. It should also be noted that neonicotinoids are known to persist and bio-accumulate in the soil [92] with their half-lives in excess of 1000 days and are capable of persisting in woody plants for over 365 days [90]. Their detections are therefore very much likely even after several months or years after application. Moreover, ever since neonicotinoids were developed from the 1980s to replace the more persistent organochlorines in the environment [91, 93], there has been a great demand for them, and particularly for imidacloprid after it was introduced to the market [94]. It was therefore not surprising that imidacloprid was found to be the most detected compound in our study. Interestingly, this observation was also confirmed by Mitchell et al [95] and Kavanagh et. al. [96]. Imidacloprid became the largest selling insecticide around the world with sales reaching $1091 million as of 2009 [97]. As a product registered and approved for 140 crops, including several crop types such as vegetables, citrus, corn, oilseed rape pome among several others, in about 120 countries [97, 98], it was therefore not unexpected that imidacloprid was detected most frequently in our included studies as it happens to have been the most used and studied compound among the neonicotinoids [99, 100].

Even though the neonicotinoids were the most detected class of pesticides residues across all studies for both cocoa and non-cocoa growing countries, our results reveal that the top three mostly detected classes of pesticides in the six cocoa growing countries were organophosphates, organochlorines and pyrethroids in that order (Fig 5 & S6 Table). Among the plausible reasons for this finding are that these pesticides have been found to be inexpensive and easily accessible and are therefore frequently used in the developing countries where most cocoa producing countries are located [101–103]. From our study, we can confirm that a total of 60 pesticides which are largely not approved for cocoa cultivation [59], were detected in studies conducted in the cocoa producing countries, the majority of which were detected in India and Mexico. The implementation of laws and regulations governing pesticides use in developing countries continue to a challenge. The ban on the use of OCPs in developed countries has witnessed remarkable successes [101] but such success has not been witnessed in the developing countries where pesticides are highly valued as a means of breaking into the global market of food production [104]. Organophosphorus pesticides (OPPs) continue to be hugely applied in the developing countries owing to their ability to inhibit disease attacks and to enhance productivity [102].

From our findings, we can confirm the presence of banned pesticides under the Stockholm Convention in three separate studies which were conducted in three different countries namely Ghana, Mexico, and India. It is however insightful to note that pollutants of organochlorine (OC) derivatives including PCBs, DTT and several other banned pesticides have been found to be persistent in the environment [105]. It is therefore not possible to confirm from our study whether detected banned pesticides were recently or previously applied. Even though research findings by [106] point to the continued use of substantial amounts of banned chemical pesticides in developing countries, it must be recognised that in some developing countries such as India, DDT which has received worldwide band, is still approved for use against mosquitoes in controlling malaria [63]. This could be a plausible reason why DDT and several derivatives were repeatedly detected in studies conducted in India (S1-S4). Again, the detection of 14 banned pesticides in a single study conducted in Mexico opens the door of speculation of recent applications. Be that as it may, the detection of 22 banned pesticides residues raises serious health concerns. This is because several diseases such as obesity, diabetes, Alzhemer’s, dementia, Pakinson’s, asthma, chronic bronchitis, autism, erectile dysfunction and many psychological disorders have been linked to exposure to banned pesticides [106].

MRLs have been set for honeybees and hive matrices including honey by the European Commission [34]. In the present study, we found that 13% of included studies detected pesticides whose concentrations exceeded allowable limits required for human consumption (Table 3), of which three cocoa producing countries accounted for a quarter of the studies where pesticides exceeded the MRL. One further important observation from our study shows that some detected pesticides residues, which exceeded the previously specified MRLs set by the EU at the time of the study, are presently below the revised MRLs that have since been implemented in the EU. This is significant as it implies that products that were previously determined to pose a risk to human health would now be assessed as not posing any unacceptable risk. The finding of pesticides exceeding MRL is significant in at least two major respects. Human exposure to levels of pesticides exceeding MRL can cause many health-related problems. The consumption of unacceptable levels of pesticides via food are known to have many implication ranging from headaches, nausea, itching and skin irritation, restlessness, dizziness, breathing complications, neurotoxicity and chronic poisoning related diseases such as cancer and in some cases death [107]. The causes of exceedances of MRLs are varied and include the use of pesticides for reasons far beyond the sphere of the intended purpose of a particular pesticide, non-compliance to specified guidelines in labels such as overdoses and spray drifts among others [108, 109]. However adherence to good agricultural practice (GAP), where specified guidelines including detailed information on labels, are followed for the safe and sustainable production of crops and livestock to maximise profit with minimal impacts to the environment, has been touted as capable of preventing excessive leaking of plant protection products into the environment [88, 110, 111]. While we do not have detailed information of agricultural practices in the areas where MRL was exceeded, it should be considered that generally a significant proportion of applied pesticides also find their way to the general ecosystem with just about 0.01% of applied pesticides reaching its target and the rest filtering into the into general ecosystem [18, 19]. Though strict enforcement of compliance to specified MRLs in honey will help promote health safety measures, a globally standardised MRLs will ensure clarity and prevent potential confusion. We observed that the concentration of malathion was evaluated using the MRL set by India and was deemed to have exceeded national MRL in India but found to be below upper limits of the MRL set by the EU. This can be a potential source of confusion for which reason standardisation across borders will be essential. It should also be considered that while the revision of MRL can impact the assessment of honey as a food product, it does not alter the assessment of pesticide contamination levels as an indicator of potential contamination of the environment surrounding the hive.

The Limit of detection and limit of quantification are of immense importance in chromatography and have regularly been used by analytical chemist in trace analysis for determining both the presence and concentrations of analytes [112, 113]. The majority of the studies reviewed used the LOD and LOQ which were usually below the MRLs set by the EU in honey which range from 0.05 mg/kg to 0.2 mg/kg [36]. This finding provides evidence that suggests that studies included in this review largely applied analytical methods with good sensitivity. It must, however, be noted that LODs for three studies were not suitable for detecting pesticide residues below EU MRLs, compromising the extent to which their results could be considered within this study. Specifically, even though no pesticide residues were detected by [114] in a study conducted in Brazil, their reported LODs and LOQs which ranged from 0.07 mg/kg to 0.25 mg/kg and 0.02 mg/kg to 0.08 mg/kg respectively were higher than EU MRLs, and so the study’s reports of no pesticides detected cannot be interpreted to mean that there were no pesticides present at concentrations with the potential to cause harm. Similarly, although pesticides were detected by [115] and [116] in studies conducted in Brazil, the LOQs achieved for the method were at concentrations so high that their findings cannot be interpreted to mean that the pesticides detected were the only ones which were a cause for concern.

Even though the scope of this research did not extend to assess the effects of pesticides on bees, the high frequency of detection of neonicotinoids in honeys as observed in our study suggest that bees could be impacted by neonicotinoids due to exposure during foraging. In our study, concentrations of 0.736 mg/kg of imidacloprid, [117]; 0.0274 mg/kg of thiacloprid [118] and 0.0202 mg/kg of thiamethoxam [119] were confirmed in Pakistan and Poland respectively. This is worth noting as neonicotinoids act to impair the nervous system targeting the nicotinic acetylcholine receptor (nAChR), an ion channel that plays a key role in nerve signalling in insects [120] leading to eventual paralysis and death [121]. Neonicotinoids are noted to affect insects with biting and sucking mouth parts if swallowed [93] of which many beneficial insects also belong to. Toxicity levels of imidaclorpid, clothianidin and thiamethoxam in the ranges of of 0.004 to 0.075 µg/bee have been established to be lethal to bees [122–124]. Although this is below the known LD_50_s for these compounds for bees [125], they are within the range of concentrations known to have sub-lethal effects. For instance bumble bees were found to reduce learning capability and their short-term memeory impaired drastically when exposed to field realistic levels to 0.0024 mg/kg of thiamathoxam [126], which is 10-fold lower concentration than what was detected by Bargańska et al. in Poland. A similar observation was also detected in a different study where the foraging and homing success of bumble bees were impacted by exposure to 0.0024 mg/kg of thiamathoxam [127]. In another study, Straub, Villamar-Bouza [128] confirmed that the survival of honeybees was reduced by 51% as well as reduced flight activities when exposed to 0.0043 mg/kg and .0011 mg/kg concentrations of thiamethoxam and clothianidin respectively. Therefore, there may be potential sub-lethal effects of the detected pesticide residues in honey on bees which deserves further consideration.

It was observed from this study that pesticide residues were detected in 80% of both commercial and raw honeys analysed in the included studies. This finding reflects a similar observation made by Mitchell, Mulhauser [95] who confirmed the presence of neonicotinoids in 75% of 198 honeys taken directly from producers in a worldwide survey of neonicotinoids in honeys. The most striking observation made, however, was that as high as 90 % of 61 studies which analysed raw honeys confirmed the presence of pesticide residues. This was found to be higher than the finding by [95] who evaluated raw honeys. However, recognizance should be taken of the fact that our findings are not limited to only neonicotinoids but covers all pesticides evaluated in raw honeys in our studied publications. Our finding highlights the high prevalence of pesticides in the general environment. Raw honeys from a broad spectrum of both natural and agricultural landscapes were assessed in the publications of interest in this study. These included raw honeys from agricultural farmlands within forest belts in Ghana [129], apiaries located 2 miles of an oilseed [130], agroclimatic zones [131], agricultural landscape with mostly intensively managed fields, forested areas and human settlements [132], unifloral and multifloral sources [133] among several others. The foraging behaviour of bees is such that they are likely to transport these substances from the environment into the hive if they come onto contact with them which may be a source of contamination at the production base. In the present study, a very small number studies evaluated the floral background of the honey, it was therefore not possible to correlate pesticide contamination to with any specific floral resources.

## Conclusion

The current state of knowledge of studies of honey contamination from pesticides approved for cocoa cultivation has been evaluated through a systematic literature review. The studies conducted over the period have been disproportionately focused on non-cocoa growing areas leaving a huge gap of knowledge of how pesticides approved for cocoa cultivation affects bee products particularly honey and, by proxy, how prevalent these pesticides abound in the localities of cocoa production. As a crop whose production hinges on the intense application of large volumes of pesticides to prevent huge losses, continuous monitoring and strict compliance of pesticide application will ensure the correct use of pesticides, thereby ensuring that pesticide residues are kept below tolerable levels. Residue analysis in honey could serve as a proxy for monitoring the extent to which pesticides are imputed for cocoa cultivation. It is recommended that cocoa producing countries are prioritised for such studies especially in the two leading cocoa producing countries namely Ivory Coast and Ghana, which account for 70% of the world cocoa but where studies were rarely conducted. These studies could form the basis for policy formulation for sustainable and effective beekeeping and pesticide application in cocoa producing countries, especially in cocoa growing landscapes.

## Supplementary information

S1 Table. Recommended active ingredients for cocoa production (DOCX)

S2 Table. Search strings (DOCX)

S3 Table. Quality Assessment

S4 Table. Dataset for the study (XLSX)

S5 Table. List of Cocoa producing countries (DOCX)

S6 Table. Classes of pesticides detected in cocoa producing countries (XLSX)

S7 Table. PRISMA checklist (DOCX)

S1 Fig to S4 Fig. Heatmaps of dectected pesticides in studied publications(DOCX)

S5 Fig. The rate of use of extraction techniques in the included studies (DOCX)

S6 Fig. PRISMA_2020_flow_diagram_updated

S8. Appendix. Description of grading system for study appraisal

## Acknowledgement

We would like to thank Helen Sheridan and Thomas Cummins, members of my Research Study Panel at University College Dublin, for the useful inputs and discussion in taking up this study. Linzi J. Thompson of University College Dublin, Elena Ziogae of Dublin City Ubiversity and Diarmuid Stokes, the College Liaison Librarian at UCD Library, are also acknowledged for their useful discussion on how to conduct systematic literature review.

## Author Contribution

**Conceptualisation:** Richard G. Boakye, Dara A. Stanley, Blanaid White

**Data curation:** Richard G. Boakye, Blanaid White

**Formal analysis**: Richard G. Boakye, Dara A. Stanley, Blanaid White

**Methodology**: Richard G. Boakye, Dara A. Stanley, Blanaid White

**Supervision:** Dara A. Stanley, Blanaid White

**Writing-original draft:** Richard G. Boakye, Dara A. Stanley, Blanaid White

**Writing-**review and editing: Richard G. Boakye, Dara A. Stanley, Blanaid White

## Notes

### Competing Interest Statement

The authors have declared no competing interest.

### Summary of Updates

Abstract updated and supplementary material uploaded. Quality assessment of included papers added and am updated search conducted

