## Supplementary figures and images for "Honey contamination from plant protection products approved for cocoa cultivation: a systematic review of existing research and methods"

### S1 Fig

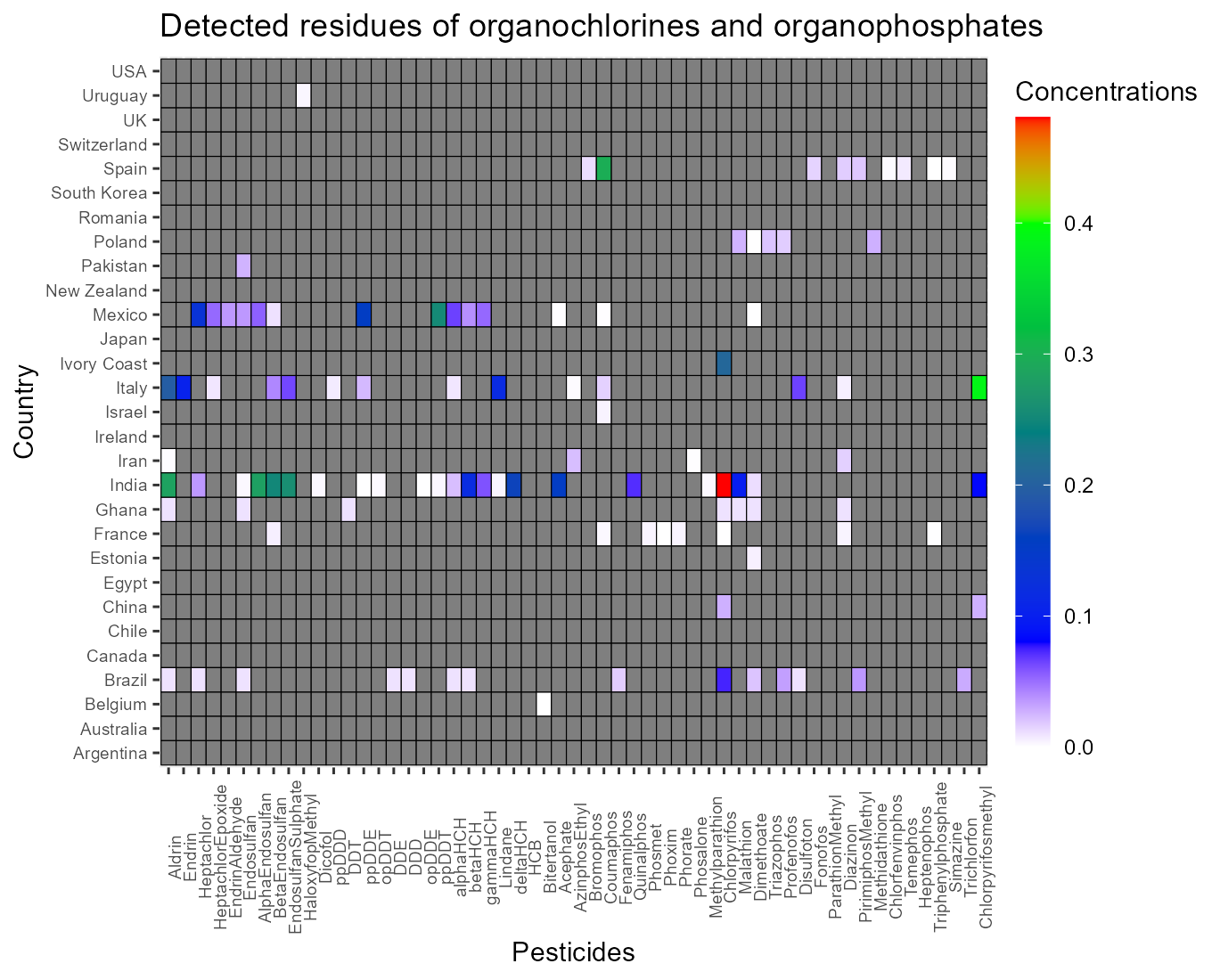

### S4 Fig

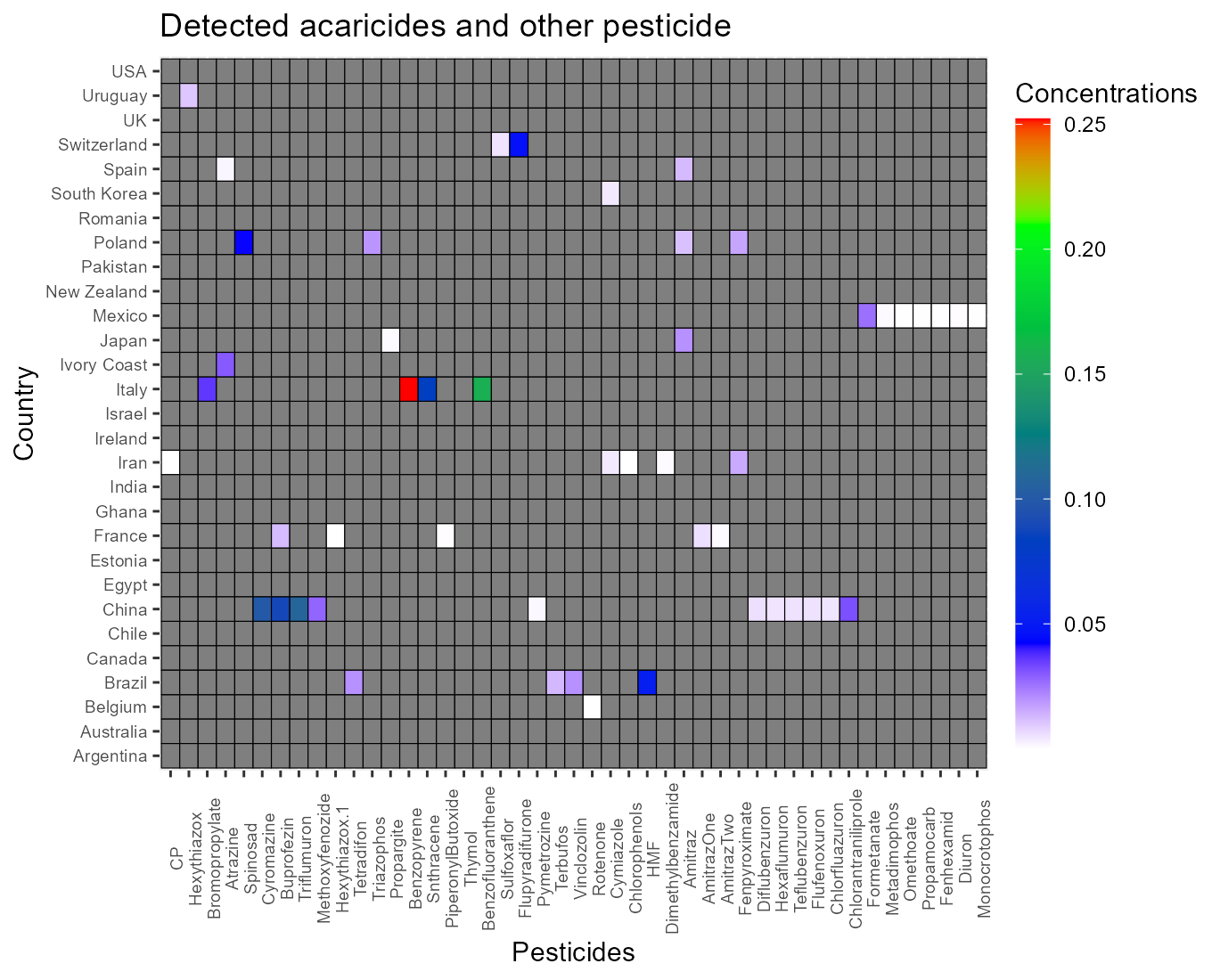

### S5 Fig

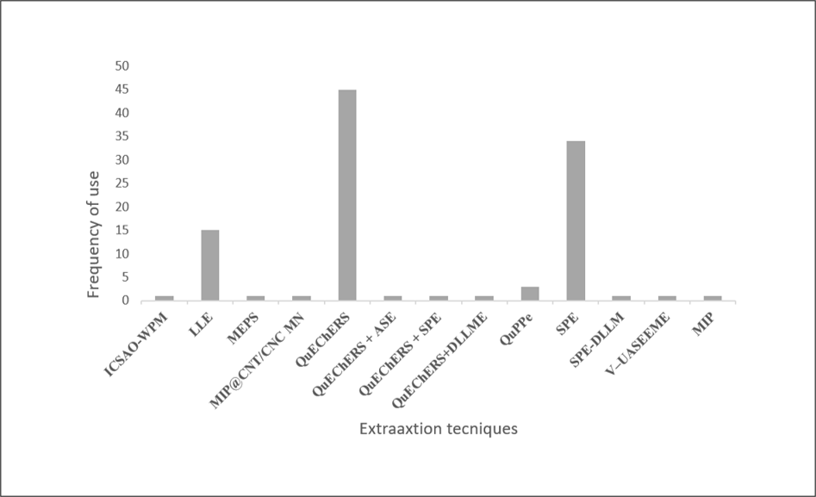

### S 2 Fig

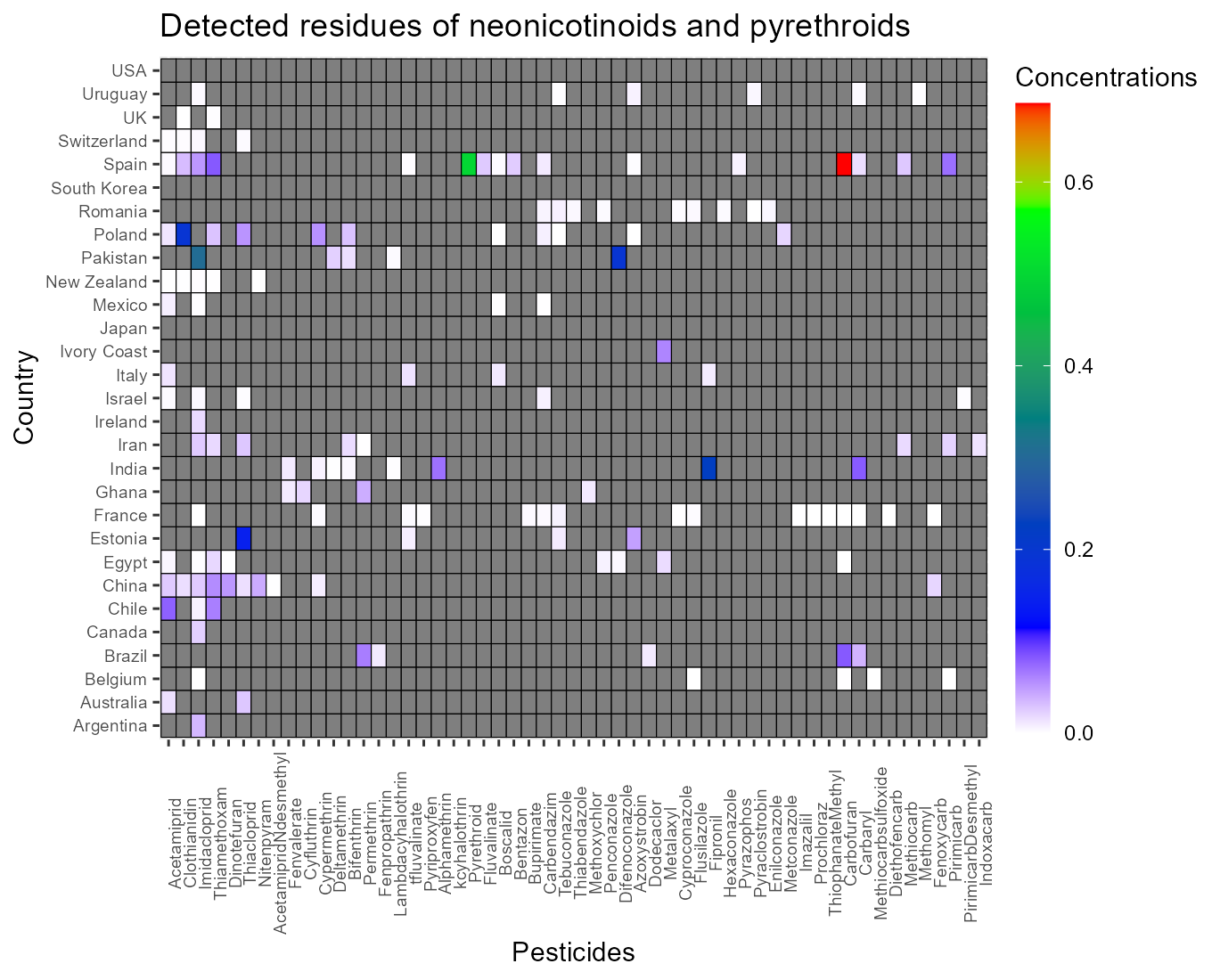

### S 3 Fig

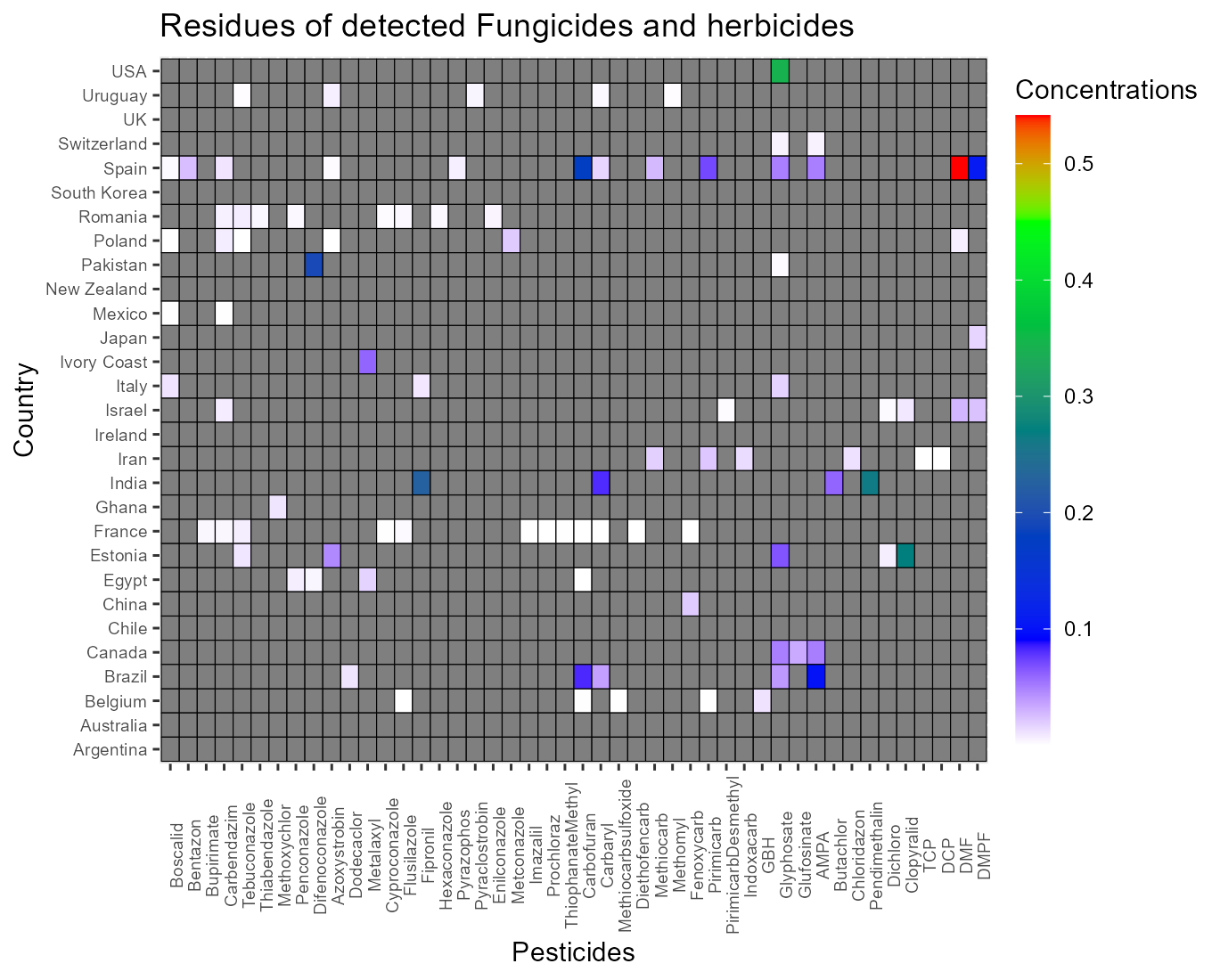
