## Supplementary material for "Honey contamination from plant protection products approved for cocoa cultivation: a systematic review of existing research and methods": S4-S6

**Heatmaps of dectected pesticides in studied publications classified by country where detected**

S1 Fig. A heat map (colour scale on the right), showing the detected organochlorines and organophosphates which were studied (x-axis) over the period and the respective countries where detections took place (y-axis). For the purpose of this figure, concentrations of each detected pesticide were averaged per country to get one value for each pesticide for each country. Units for all detected pesticides were standardised by converting to mg/kg which is the unit used by the European Union and which was adopted for this study as most of the studies compared detections to the maximum residues limits (MRL) set by the EU.

S2 Fig. A heat map (colour scale on the right), showing the neonicotinoids, pyrethroids and carbamates which were studied (x-axis) over the period and the respective countries where detections took place (y-axis). For the purpose of this figure, concentrations of each detected pesticide were averaged per country to get one value for each pesticide for each country. Units for all detected pesticides were standardised by converting to mg/kg which is the unit used by the European Union and which was adopted for this study as the majority of the studies compared detections to the maximum residues limits (MRL) set by the EU

S3 Fig. A heat map (colour scale on the right), showing the fungicides and herbicides which were studied (x-axis) over the period and the respective countries where detections took place (y-axis). For the purpose of this figure, concentrations of each detected pesticide were averaged per country to get one value for each pesticide for each country. Units for all detected pesticides were standardised by converting to mg/kg which is the unit used by the European Union and which was adopted for this study as majority of the studies compared detections to the maximum residues limits (MRL) set by the EU

Fig S4. A heat map (colour scale on the right), showing the acaricides and other pesticides which were studied (x-axis) over the period and the respective countries where detections took place (y-axis). For the purpose of this figure, concentrations of each detected pesticide were averaged per country to get one value for each pesticide for each country. Units for all detected pesticides were standardised by converting to mg/kg which is the unit used by the European Union and which was adopted for this study as majority of the studies compared detections to the maximum residues limits (MRL) set by the EU

**S5 Fig. The rate of use of extraction techniques in the included studies. The Quick, Easy, Cheap, Effective, Rugged and Safe technique popularly called QuEChERS was the most dominant extraction techniques having been applied in 37 of the included studies.**
