## Supplementary material for "Honey contamination from plant protection products approved for cocoa cultivation: a systematic review of existing research and methods": S1 Table

**S1 Table. Summary of approved active ingredients for cocoa production. These pesticides are recommended for cocoa cultivation in major cocoa producing countries in West Africa which account for 70% of the World’s cocoa.**

| **Insecticides (active) ingredients** | **Fungicides** | **Herbicides** | **Sources** |
| --- | --- | --- | --- |
| Cypermethrin, Capsaicin, Imidacloprid, Chlorpyriphos, Dimethoate, Deltamethrin, Thiamethoxam, Acetaprimid, Bifenthrin, Pyrethrum, Alpha-cypermethrin, Teflubenzuron, Lambda cyhalothrin, Indoxacarb,  Chlorantraniliprole, Fipronil, Sulfoxaflor, Etofenprox,  Pirimiphosmethyl and Promecarb | Copper oxide, Metalaxyl, Mancozeb, Maned, Benalaxyl, Benomyl, Copper hydroxide, Metalaxyl-M, Copper II hydroxide, Mefenoxam, Cupper (I) oxide, Dicopper chloride trihydroxide, Dimethomorph, Fluazinam, Cuprous hydroxide and  Cupric hydroxide | Glyphosate  Paraquat | (Mahob et al., 2014); (Bateman, 2010); (Boamah, 2020);  (Ninsin and Adu-Acheampong, 2017); (Antwi-Agyakwa et al., 2015); (Afrane and Ntiamoah, 2011)  (Oyekunle et al., 2017)  (Mokwunye et al., 2014) |
