## Supplementary material for "Honey contamination from plant protection products approved for cocoa cultivation: a systematic review of existing research and methods": S4 Table

**S2 Table. Search strings used to retrieve articles from the search engines.**

| **Search strings** | **Search engine** | **Targeted category** |
| --- | --- | --- |
| (cypermethrin OR capsaicin OR chlorpyriphos OR imidacloprid OR dimethoate OR deltamethrin OR thiamethoxam OR acetaprimid) AND (honey)  (bifenthrin OR pyrethrum OR alpha-cypermethrin OR teflubenzuron OR “lambda cyhalothrin” OR indoxacarb OR chlorantraniliprole OR fipronil OR sulfoxaflor OR etofenprox OR pirimiphosmethyl OR promecarb) AND (honey) | Web of Science Core Collection, PubMed, and Scopus | Insecticides |
| (“copper oxide” OR metalaxyl OR “cuprous hydroxide” OR mancozeb OR maned OR benalaxyl OR benomyl OR “Copper hydroxide” OR Metalaxyl-M OR “copper II hydroxide” OR mefenoxam OR “cupper (I) oxide” OR “dicopper chloride trihydroxide” OR dimethomorph OR fluazinam OR “cuprous hydroxide” OR “cupric hydroxide”) AND (honey) | Web of Science Core Collection, PubMed, and Scopus | Fungicides |
| (glyphosate OR paraquat) AND (honey) | Web of Science Core Collection, PubMed, and Scopus | Herbicides |
