## Supplementary material for "Honey contamination from plant protection products approved for cocoa cultivation: a systematic review of existing research and methods": S5 Table

**S3 Table: Cocoa producing countries according to metric tons of cocoa beans produced . Studies were conducted only in eight cocoa-producing countries including one each in the top two countries**

| **Ranking** | **Country** | **Cocoa Production (metric tons)** | **Pop. 2021** |
| --- | --- | --- | --- |
| 1 | Ivory Coast | 2034000 | 27053.629 |
| 2 | Ghana | 883652 | 31732.129 |
| 3 | Indonesia | 659776 | 276361.783 |
| 4 | Nigeria | 328263 | 211400.708 |
| 5 | Cameroon | 295028 | 27224.265 |
| 6 | Brazil | 235809 | 213993.437 |
| 7 | Ecuador | 205955 | 17888.475 |
| 8 | Peru | 121825 | 33359.418 |
| 9 | Dominican Republic | 86599 | 10953.703 |
| 10 | Colombia | 56808 | 51265.844 |
| 11 | Papua New Guinea | 44504 | 9119.01 |
| 12 | Uganda | 31312 | 47123.531 |
| 13 | Mexico | 27287 | 130262.216 |
| 14 | Venezuela | 23349 | 28704.954 |
| 15 | Togo | 22522 | 8478.25 |
| 16 | India | 19000 | 1393409.038 |
| 17 | Sierra Leone | 14670 | 8141.343 |
| 18 | Haiti | 14173 | 11541.685 |
| 19 | Guatemala | 11803 | 18249.86 |
| 20 | Madagascar | 11010 | 28427.328 |
| 21 | Guinea | 10638 | 13497.244 |
| 22 | Liberia | 8552 | 5180.203 |
| 23 | Tanzania | 8548 | 61498.437 |
| 24 | Philippines | 7009 | 111046.913 |
| 25 | Nicaragua | 6600 | 6702.385 |
| 26 | Bolivia | 5518 | 11832.94 |
| 27 | Solomon Islands | 4940 | 703.996 |
| 28 | Republic of the Congo | 4000 | 5657.013 |
| 29 | DR Congo | 3758 | 92377.993 |
| 30 | Sao Tome and Principe | 2778 | 223.368 |
| 31 | Vanuatu | 1813 | 314.464 |
| 32 | Sri Lanka | 1291 | 21497.31 |
| 33 | Malaysia | 1029 | 32776.194 |
| 34 | Grenada | 800 | 113.021 |
| 35 | Honduras | 751 | 10062.991 |
| 36 | Panama | 662 | 4381.579 |
| 37 | Costa Rica | 545 | 5139.052 |
| 38 | Samoa | 479 | 200.149 |
| 39 | Angola | 442 | 33933.61 |
| 40 | Guyana | 429 | 790.326 |
| 41 | Equatorial Guinea | 413 | 1449.896 |
| 42 | El Salvador | 357 | 6518.499 |
| 43 | Trinidad and Tobago | 320 | 1403.375 |
| 44 | Dominica | 312 | 72.167 |
| 45 | Jamaica | 305 | 2973.463 |
| 46 | Belize | 244 | 404.914 |
| 47 | Cuba | 231 | 11317.505 |
| 48 | Saint Vincent and the Grenadines | 222 | 111.263 |
| 49 | Gabon | 187 | 2278.825 |
| 50 | Timor-Leste | 176 | 1343.873 |
| 51 | Central African Republic | 149 | 4919.981 |
| 52 | Thailand | 125 | 69950.85 |
| 53 | Saint Lucia | 51 | 184.4 |
| 54 | Comoros | 40 | 888.451 |
| 55 | Micronesia | 33 | 116.254 |
| 56 | Fiji | 12 | 902.906 |
| 57 | Suriname | 5 | 591.8 |
| 58 | American Samoa | 1 | 55.1 |

**Sourced at** World Population Review [**https://worldpopulationreview.com/country-rankings/cocoa-producing-countries**](https://worldpopulationreview.com/country-rankings/cocoa-producing-countries) **on 10/12/2021**
