## Supplementary material for "Honey contamination from plant protection products approved for cocoa cultivation: a systematic review of existing research and methods": S8 Appendix

**Appendix :** **Description of scoring scheme for included studies (adapted from [1]).**

**Scoring:** “yes”=2; “partial”=1 & “no”=0, “NA”=not applicable and not included in total score

**How to summarise the total score per study**

Total score = (number of “yes” scored “yes” * 2) + (number of “partials” * 1)

Summary score =total score/total possible score

(There 11 criteria/questions, total possible score is therefore 22)

**Description of appraisal questions**

**1. Is the study objective clearly stated?**

**Yes:** Easily identified in the abstract and in the introduction.

**Partial:** Study objective not specific or incompletely described. Or where the whole paper must be read before understanding the aim of a study will amount to study objective not being sufficiently described

**No:** Study objective not stated or inexplicable

**2.0 Is the study design sufficiently described?**

**Yes:** Study design clearly described in the method section and appropriate to answer the study objective

**Partial:** Study design not clearly identified or stated but not appropriate to answer the study aim

**No**: Study design not stated or cannot be identified

**3.0 Are all honey samples from one specific country?**

**Yes:** All samples taken from only one country

**No.** Where honey samples are from multiple countries or where the country name is not state (where honey samples are not from one country study cannot be assigned to any specific country and is excluded for further analysis)

**4.0 If pesticide residues were detected in honey, were the concentrations quantified in honey?**

**Yes:** Concentrations of pesticide residues quantified and reported. This provides opportunity for further analysis

**Partial:** If detected pesticides are not quantified. Detections of pesticide residues were reported but their concentrations were not reported in results or abstract.

**No:** Where only blank honey is analysed but no actual concentrations of pesticides necessarily quantified in real honey. Where no concentrations of pesticides are quantified in actual honey, the study is excluded for further analysis.

NA: Not applicable is entered if pesticide residues were not detected. Where no detections were made and therefore no quantification required, scoring for this response is not included in the total sum of scoring

**5.0 Is the limit of detection (LOD) below the maximum residues (MRL) of studied compounds?**

**Yes.** If all LODs of detected pesticides residues are below the specified MRL. On this occasion the analytical technique is deemed sufficiently sensitive to detect compounds being studied

**Partial**: Where study reports detection but does not give LOD

**Partial**: If some LOD is below with other being higher the EU MRL

**No:** If all LODs are higher than EU MRL

**6.0** Is the limit of quantification (LOQ) below the maximum residue limit (MRL).

**Yes**. If all LODs of detected pesticide residues are below the specified EU MRL. On this occasion the analytical technique is deemed sufficiently sensitive to quantify compounds being studied

**Partial:** If some LOQ is below with others being higher the MRL

**No**: If all LODs are higher than MRL

**7.0 Is the extraction technique clearly described /appropriate?**

**Yes:** Extraction technique clearly defined. Very appropriate based on scientifically acceptable or standard procedures.

**Partial:** Vaguely defined or incompletely described. Or where the whole paper must be read before understanding procedure applied. Or where extraction technique is not applied to analyse residues of pesticides in actual honey.

**No:** Where extraction procedure is not described or not based on standard procedures.

**8.0 Is the analytical technique clearly described/appropriate?**

**Yes**: Analytical technique clearly defined. Very appropriate based on scientifically acceptable or standard procedures

**Partial:** Vaguely defined or incompletely described. Or where the whole paper must be read before understanding procedure applied

**No:** Where analytical technique is not described or not based on standard procedures

**9.0 Is the type of source of honey (raw or commercial) indicated?**

**Yes:** If the honey is indicated as being either raw honey or commercial honey

**No:** if the types of honey samples analysed is not indicated

**10.0 Are study results reported in sufficient details?**

**Yes:** All major outcomes and any secondary outcomes are reported

**Partial:** Quantitative results reported for some outcomes. Or difficult to assess as study question/objective not fully described even though results may seem appropriate

**No:** Results reported for some samples or pesticide studied or reported proportion does not account for the entire study sample.

**11.0 Are the conclusions supported by results?**

**Yes**: All the conclusions are supported by data. Conclusions based on results relevant to the study questions whether negative or positive

**Partial:** Some of major conclusions are supported by the data but others are not.

**No:** Either no or very small minority of the major conclusions are supported by the data. Or conclusions missing
